# Early anteroposterior regionalisation of human neural crest is shaped by a pro-mesodermal factor

**DOI:** 10.1101/2021.09.24.461516

**Authors:** Antigoni Gogolou, Celine Souilhol, Ilaria Granata, Filip J Wymeersch, Ichcha Manipur, Matthew Wind, Thomas JR Frith, Maria Guarini, Alessandro Bertero, Christoph Bock, Florian Halbritter, Minoru Takasato, Mario R Guarracino, Anestis Tsakiridis

## Abstract

The neural crest (NC) is an important multipotent embryonic cell population and its impaired specification leads to various developmental defects, often in an anteroposterior (A-P) axial level-specific manner. The mechanisms underlying the correct A-P regionalisation of human NC cells remain elusive. Recent studies have indicated that trunk NC cells, the presumed precursors of the childhood tumour neuroblastoma, are derived from neuromesodermal-potent progenitors of the postcranial body (NMPs). Here we employ human embryonic stem cell differentiation to define how NMP-derived NC cells acquire a posterior axial identity. We show that TBXT, a pro-mesodermal transcription factor, mediates early posterior NC regionalisation together with WNT signalling effectors. This occurs by TBXT-driven chromatin remodelling via its binding in key enhancers within HOX gene clusters and other posterior regulator-associated loci. In contrast, posteriorisation of NMP-derived spinal cord cells is TBXT/WNT-independent and takes place under the influence of FGF signalling. Our work reveals a previously unknown role of TBXT in influencing posterior NC fate and points to the existence of temporally discrete, cell type-dependent modes of posterior axial identity control.

## Introduction

The neural crest (NC) is a multipotent cell population, which arises in the dorsal neural plate/non-neural ectoderm border region during vertebrate embryogenesis and generates a variety of cell types following epithelial-to-mesenchymal transition and migration through diverse routes. NC cells emerge at all levels of the anteroposterior (A-P) axis and their A-P position determines the identity of their derivatives: cranial NC produces mesoectodermal cell types (e.g. dermis, cartilage, bone), melanocytes, neurons and glia colonizing the head; vagal NC cells, arising between somites 1–7, contribute (together with their sacral counterparts emerging at axial levels posterior to somite 28) predominantly to the enteric nervous system and includes a subpopulation (cardiac NC) that generates various heart structures; trunk NC generates dorsal root/sympathetic neurons, adrenal chromaffin cells and melanocytes (reviewed in (Le Douarin et al., 2004, Rocha et al., 2020, Rothstein et al., 2018). Defects in the specification, differentiation or migration of NC cells lead to a wide spectrum of serious developmental disorders, often in an axial level-specific manner and are known collectively as neurocristopathies (Pilon, 2021, Vega-Lopez et al., 2018). The use of human pluripotent stem cell (hPSC) differentiation offers an attractive platform for the study of human NC biology and neurocristopathies as well as an indefinite *in vitro* source of clinically relevant NC-associated cell populations for regenerative medicine applications. However, the cellular and molecular mechanisms directing the acquisition of distinct A-P axial identities by human NC cells remain largely undefined. In turn, this obviates the design of optimised hPSC differentiation protocols aiming to produce NC derivatives as well as the dissection of the links between human neurocristopathy emergence and the axial identity of the NC cells affected.

*In vivo*, the A-P patterning of the vertebrate embryonic body and its nascent cellular components relies on the coordinated action of Hox gene family members (arranged as paralogous groups (PG) in four distinct chromosomal clusters: A, B, C and D). In mammals, transcriptional activation of Hox genes is initiated during gastrulation, within the posterior part of the embryo around the primitive streak and proceeds in a sequential manner reflecting their 3’-to-5’ genomic order, a phenomenon described as temporal collinearity or the Hox clock (Deschamps & Duboule, 2017, Dolle et al., 1989, Izpisua-Belmonte et al., 1991). The Hox clock continues to operate after gastrulation and until the end of somitogenesis, within a posterior growth zone located in/around the caudal lateral epiblast-late primitive streak and later the tail bud (Deschamps & Duboule, 2017, Wymeersch et al., 2019, Wymeersch et al., 2021). The colinear activation of Hox genes within these posterior regions is thought to be tightly coupled to the assignment of the terminal A-P coordinates of the cell lineages that make up the postcranial axis, including the NC (Deschamps & Duboule, 2017, Wymeersch et al., 2019, Wymeersch et al., 2021). Eventually, the terminal expression domains of Hox genes along the A-P axis corresponds to their ordering within their chromosomal clusters so that the earliest activated, 3’ Hox PG members are expressed more anteriorly compared to their later-activated 5’ counterparts. In the case of NC, anterior cranial NC is Hox-negative, posterior cranial NC cells express Hox PG(1-3) genes, while vagal NC cells are marked by the expression of Hox PG(3-5) members and positivity for Hox PG(5-9) genes denotes a trunk NC character (Rocha et al., 2020).

The post-gastrulation posterior growth region is marked by high levels of Wnt/Fgf signalling activity and harbours a pool of multipotent axial progenitors driving embryonic axis elongation in a head-tail direction. These include neuromesodermal progenitors (NMPs) that generate a large fraction of the spinal cord as well as presomitic/paraxial mesoderm, the building blocks of the musculoskeleton (reviewed in (Henrique et al., 2015, Wymeersch et al., 2021). NMPs are marked by the co-expression of neural and mesodermal genes, such as those encoding the transcription factors *Sox2* and *Brachyury (T)* (Guillot et al., 2021, Henrique et al., 2015, Martin & Kimelman, 2012, Olivera-Martinez et al., 2012, Tsakiridis et al., 2014), as well as a number of other posteriorly expressed genes such as *Nkx1-2*, *Cdx2*, *Tbx6* and *Hox* family members (Amin et al., 2016, Dias et al., 2020, Gouti et al., 2017, Guillot et al., 2021, Javali et al., 2017, Koch et al., 2017, Rodrigo Albors et al., 2018, Wymeersch et al., 2019, Young et al., 2009).

Fate mapping, lineage tracing and single cell transcriptomics studies in vertebrate embryos have revealed that trunk NC cells, which give rise to neuroblastoma (the most common extracranial solid tumour in infants) following oncogenic transformation, are derived from NMPs (Javali et al., 2017, Lukoseviciute, 2021, Shaker et al., 2021, Tzouanacou et al., 2009, Wymeersch et al., 2016). Moreover, recent work has demonstrated that the *in vitro* generation of trunk NC cells and their derivatives from hPSCs relies on the induction of a NMP-like intermediate following stimulation of WNT/FGF signalling pathways (Abu-Bonsrah et al., 2018, Cooper, 2020, Faustino Martins et al., 2020, Frith et al., 2018, Frith & Tsakiridis, 2019, Gomez, 2019, Hackland et al., 2019, Kirino et al., 2018). These and other studies have pointed to a model where a generic posterior axial identity is installed early within the NMP precursors of trunk NC cells under the influence of WNT/FGF activities prior to their differentiation and commitment to a NC fate (Frith et al., 2018, Metzis et al., 2018). However, it is unclear whether induction of a developmentally plastic, Brachyury-positive state is an obligatory step in the ‘posteriorisation’ of prospective NC cells or trunk NC cells can still acquire a posterior axial identity without passing through a mesoderm-competent progenitor stage.

Here we dissect the links between human NMP induction and trunk NC specification using hPSC differentiation as a model. We show that disruption of NMP ontogeny via knockdown of the NMP/mesoderm regulator TBXT (the human homologue of Brachyury) abolishes the acquisition of a posterior axial identity by NC cells without affecting adoption of an NC fate. This is linked to a failure of WNT-FGF-treated hPSCs to activate properly the expression of HOX genes. We demonstrate that TBXT mediates early posteriorisation by directly orchestrating an open chromatin landscape in HOX clusters and key WNT signalling-linked loci. In contrast, control of trunk HOX gene expression/posteriorisation in NMP-derived early central nervous system (CNS) spinal cord progenitors appears to be TBXT/WNT-independent and programmed after NMP differentiation primarily under the action of FGF signalling. Collectively, our data reveal a previously unappreciated role for TBXT in prospectively shaping the A-P patterning of NC cells. They also indicate the existence of two distinct phases of posterior axial identity control: (i) an early NMP-based that involves the TBXT/WNT-driven establishment/fixing of a HOX-positive, posterior character in uncommitted progenitors followed by its “transmission” to their downstream trunk NC derivatives and, (ii) a later one, which is based on the FGF-driven sculpting of a posterior axial identity in cells transiting toward a spinal cord neural fate, post-NMP differentiation.

## Results

### TBXT controls posterior axial identity acquisition by NMP-derived neural crest cells

We have previously described the *in vitro* generation of human neuromesodermal (NM)-potent cell populations resembling embryonic NMPs, following a 3-day treatment of hPSCs with the WNT agonist CHIR99021 (CHIR) and FGF2 (Frith et al., 2018, Gouti et al., 2014). We showed that TBXT^+^ hPSC-derived NMPs give rise to TBXT-negative early trunk NC cells (marked by the expression of HOX PG(1-9) members/CDX2 together with NC markers such as SOX10) following re-plating and further 5-day culture in the presence of CHIR and controlled levels of BMP signalling (Frith et al., 2018, Frith & Tsakiridis, 2019). We hypothesised that if acquisition of a posterior axial identity by hPSC-derived trunk NC cells relies on the induction of NM-potent progenitors then early disruption of NMP induction/mesoderm formation competence should impair subsequent NC posteriorisation. To test this, we examined the effects of attenuating TBXT, a well-established regulator of NMP maintenance and mesodermal differentiation (Amin et al., 2016, Gouti et al., 2014, Guibentif et al., 2021, Koch et al., 2017, Martin & Kimelman, 2010, Rashbass et al., 1991) using a human embryonic stem cell (hESC) line engineered to exhibit shRNA-mediated, tetracycline (Tet)-inducible knockdown of *TBXT* expression (Bertero et al., 2016). We confirmed that Tet addition during the differentiation of these hESCs toward NMPs induced a considerable reduction in TBXT expression both at the transcript and protein level compared to untreated controls (**Fig. 1A-D**). No effect on TBXT induction was observed in NMPs generated from an isogenic control Tet-inducible shRNA hESC line targeting the *B2M* gene (Bertero et al., 2016) following Tet treatment (**Fig. EV1A**).

**Figure 1.**
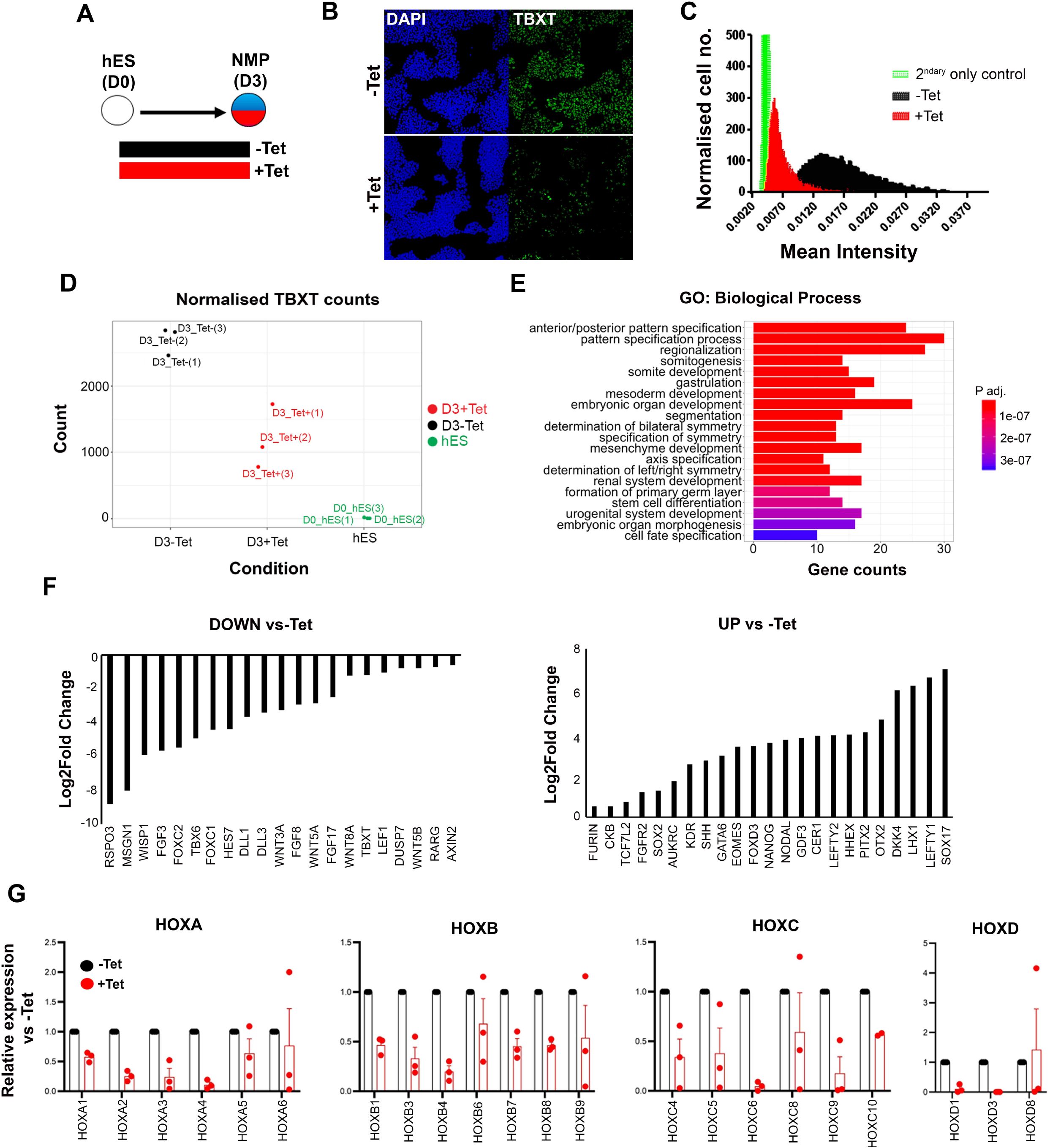
Effect of TBXT reduction on hPSC-derived NMPs. **(A)** Differentiation/treatment scheme. **(B)** Immunofluorescence analysis of the expression of TBXT in shRNA hESC-derived NMPs in the presence and absence of tetracycline (Tet). **(C)** Mean fluorescence intensity of TBXT protein in Tet-treated and control NMP cultures. **(D)** Normalised expression values of TBXT transcripts in control, Tet-treated NMPs and undifferentiated hES cell samples following RNA-seq analysis. **(E)** GO term analysis of differentially expressed genes in hESC-derived NMPs following TBXT knockdown. **(F)** Representative significantly down- and upregulated transcripts following TBXT depletion. **(G)** qPCR expression analysis of indicated HOX genes in control vs Tet-treated NMPs.

To define the impact of TBXT depletion during the transition of hPSCs toward an NMP state, we carried out transcriptome analysis of Tet-treated and control hESCs cultured in NMP-inducing conditions (i.e. presence of CHIR and FGF2) for 3 days, using RNA sequencing (RNA-seq). We found that 346 and 293 genes were significantly up- and down-regulated respectively in TBXT knockdown cells compared to untreated controls (P adj<0.05, Wald test; log2FC > |0.5|)) (**Appendix Table S1**). Gene Ontology (GO) biological processes enrichment analysis revealed that most differentially expressed genes were established regulators of A-P regionalisation/posterior development (**Fig. 1E, Appendix Table S2**). Significantly downregulated genes included presomitic mesoderm-associated transcription factors (*TBX6*, *MSGN1*, *FOXC1/2*) as well as WNT (e.g. *RSPO3*, *WISP1*, *WNT5A/B*, *WNT8A*, *LEF1*), FGF (FGF3/8/17, *DUSP7*) and NOTCH (*HES7*, *DLL1/3*) signalling pathway-linked transcripts (**Fig. 1F, Appendix Table S1)**, most of which have been reported to be present in NMPs and their immediate mesodermal derivatives *in vivo* (Guillot et al., 2021, Koch et al., 2017, Wymeersch et al., 2019). Moreover, we found that TBXT knockdown triggered a reduction in the expression of various HOX genes belonging to anterior, central and posterior PG(1-9) (**Fig. 1G, Appendix Table S1**) while no effect was observed in Tet-treated *B2M* shRNA hESC-derived NMPs (**Fig. EV1B**) ruling out the possibility that the observed decrease in HOX transcript levels may be due to nonspecific effects of Tet and/or interference of shRNAs with the micro RNA processing machinery. On the contrary, the most-highly upregulated genes included anterior visceral endoderm (AVE)/endoderm, anterior neurectoderm and pluripotency/early post-implantation epiblast-associated genes such as *SOX17*, *LEFTY1/2*, *OTX2*, *CER1*, *HHEX*, *NANOG* and *GDF3* (**Fig. 1F, Appendix Table S1)**. Taken together, these results indicate that TBXT knockdown severely impairs the induction of NMPs and their immediate presomitic mesoderm progenitor derivatives from hPSCs. They also suggest that TBXT directs the establishment of a “posteriorising” signalling environment associated with early HOX gene activation, upon pluripotency exit, since in its absence hPSCs adopt an identity that resembles the anterior epiblast despite the combined presence of caudalising WNT (CHIR) and FGF (FGF2) signalling agonists.

We next assessed the effect of TBXT disruption/failure to induce NMPs/presomitic mesoderm on trunk NC specification. To this end, we attempted to generate trunk NC cells from TBXT inducible shRNA hESCs via an NMP intermediate, treating cells initially with CHIR-FGF2 for 3 days to induce NMPs, followed by their re-plating under NC-promoting conditions (i.e. low CHIR and moderate BMP levels) for a further 5 days, as previously described (Frith et al., 2018, Frith & Tsakiridis, 2019), and either in the continuous presence or absence of Tet (**Fig. 2A**). We found that the expression of most HOX PG(1-9) genes examined was dramatically reduced in the resulting Tet-treated cultures compared to their untreated counterparts (**Fig. 2B, C**) indicating that failure of TBXT-depleted NMPs to induce properly HOX gene expression persists in their NC derivatives. The expression levels of *CDX2*, an early trunk NC regulator (Sanchez-Ferras et al., 2016, Sanchez-Ferras et al., 2012) were also found to be severely decreased (**Fig. 2D**). A modest but statistically significant reduction in the levels some NC-associated transcripts such as *SOX10* and *PAX3* was observed in the presence of Tet (P value<0.05 and <0.01 respectively; paired t-test) (**Fig. 2D**) though no obvious changes in SOX10 protein levels were detected (**Fig. 2E**). We also observed a trend towards a slight upregulation in the expression of the anterior neural crest markers *OTX2* and *ETS1* (Simoes-Costa & Bronner, 2016) as well as the neural progenitor marker *SOX1* (Wood & Episkopou, 1999) following TBXT knockdown (**Fig. 2D**). Based on these data we conclude that the acquisition of a posterior axial identity by trunk NC cells occurs at an early stage, prior to the emergence of SOX10^+^ definitive NC cells from NMPs. Intriguingly, this process is shaped by the action of TBXT, an NMP/pro-mesodermal regulator.

**Figure 2.**
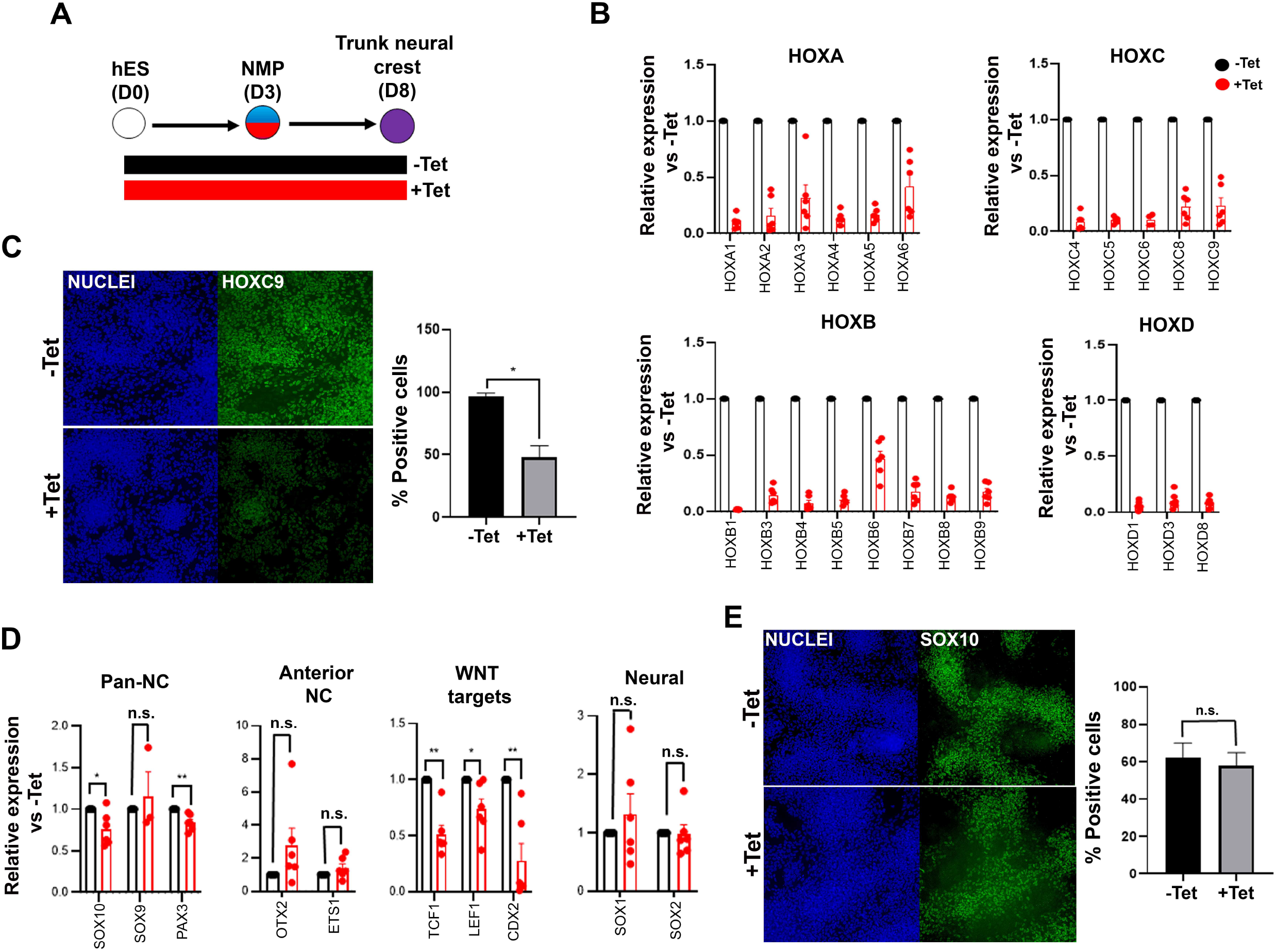
TBXT depletion impairs posterior axial identity acquisition by neural crest. **(A)** Differentiation/treatment scheme. **(B)** qPCR expression analysis of indicated HOX genes in control vs Tet-treated NMP-derived trunk neural crest cells. **(C)** Immunofluorescence analysis of the expression of HOXC9 in control vs Tet-treated NMP-derived trunk neural crest cells. Quantification of HOXC9^+^ cells in the presence and absence of Tet is also shown. **(D)** qPCR expression analysis of indicated markers in control vs Tet-treated NMP-derived trunk neural crest cells. **(E)** Immunofluorescence analysis of the expression of SOX10 in control vs Tet-treated NMP-derived trunk neural crest cells. Quantification of SOX10^+^ cells in the presence and absence of Tet is also shown. NC, neural crest. *P<0.05, **P<0.01, n.s. not significant (Paired t-test).

### Early encoding of a posterior axial identity in NMP-derived neural crest cells is primarily WNT-dependent

Our RNA-seq data revealed the concomitant downregulation of WNT targets and upregulation of WNT antagonists (e.g. *DKK4*) in day 3 FGF-CHIR-treated cultures in the presence of Tet (**Appendix Table S1**, **Fig. 1F)** suggesting that the inability of TBXT-depleted NC cells to activate HOX genes may be linked to an early reduction in WNT activity. This is further supported by our observation that TBXT knockdown-triggered reduction of HOX gene expression in our NC cultures was also accompanied by a significant reduction in the expression of WNT signalling target genes such as *TCF1*, *LEF1* and *CDX2* (**Fig. 2D**). We therefore sought to determine the temporal effects of WNT signalling on posterior axial identity acquisition during the transition from hPSCs to NMPs and subsequently NC. We first assessed the effects of WNT signalling inhibition during the 3-day induction of NMPs from hPSCs, by removing CHIR and treating the cultures with the tankyrase inhibitor XAV939 (XAV) (Huang et al., 2009) in order to eliminate any endogenous paracrine signalling, in the presence of FGF2 (the other major signalling agonist included in the media) (**Fig. 3A**). Given the reported role of FGF signalling in controlling the HOX clock in NMPs and their derivatives (Delfino-Machin et al., 2005, Hackland et al., 2019, Liu et al., 2001, Mouilleau et al., 2021, Nordstrom et al., 2006), we also tested the effect of attenuating this pathway in a similar manner, by omitting FGF2 and culturing cells only in the presence of CHIR and the FGF pathway-MEK1/2 inhibitor PD0325901 (PD03) between days 0-3 (**Fig. 3A**). Signalling inhibition was verified by confirming the downregulation/extinction of WNT (*AXIN2*, *TCF1*, *LEF1*) and FGF signalling target genes (*SPRY4*) in inhibitor-treated day 3 cultures relative to untreated controls (**Fig. 3B**). Our results confirmed that maximal induction of NMP markers (*TBXT*, *CDX2* and *NKX1-2*) and all HOX genes examined can be achieved only in the combined presence of WNT and FGF agonists (**Fig. 3C, 3D**, white bars). However, WNT signalling stimulation alone, in the absence of any FGF activity, rescued the induction of most HOX genes and *TBXT/CDX2* with variable efficiency (**Fig. 3D**, red bars) indicating that WNT is the main instructive pathway controlling the HOX clock and expression of major axis elongation regulators during the differentiation of hPSCs toward NMPs. HOXC genes were an exception, as their activation was found to be equally dependent on WNT and FGF signalling (**Fig. 3D**) suggesting the additional existence of HOX cluster-specific modes of transcriptional control, in line with similar findings in the embryo (Ahn et al., 2014, van den Akker et al., 2001).

**Figure 3.**
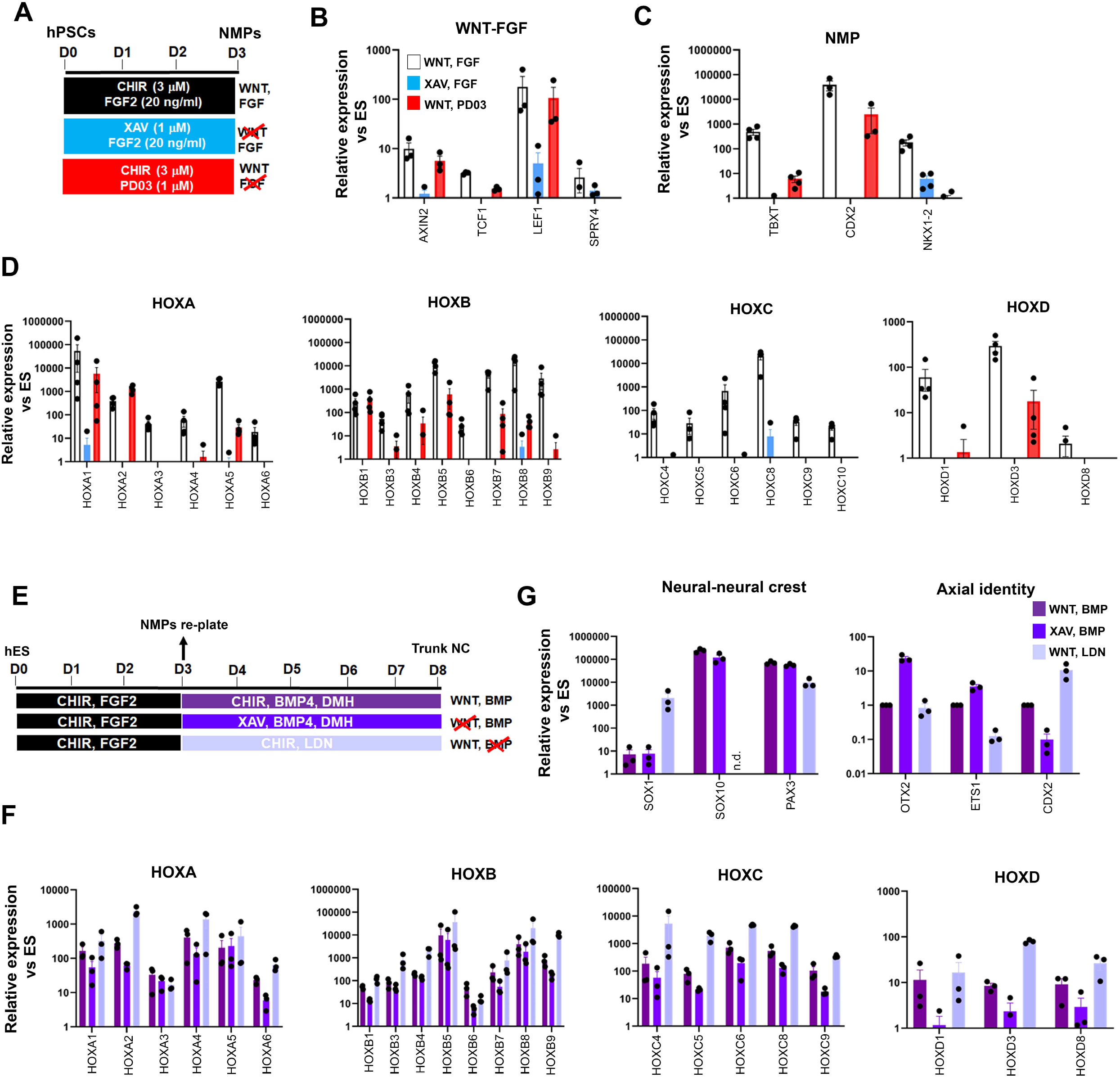
Early programming of a posterior axial identity in NMP-derived neural crest cells is primarily WNT-dependent. **(A)** Scheme of treatments during the differentiation of hPSCs toward NMPs. **(B)** qPCR expression analysis of representative WNT-FGF targets in NMP cultures treated with the indicated combinations of WNT-FGF agonists/antagonists. **(C-D)** qPCR expression analysis of key NMP markers **(C)** and HOX genes **(D)** in NMP cultures treated with the indicated combinations of WNT-FGF agonists/antagonists. **(E)** Scheme of treatments during the differentiation of hPSC-derived NMPs toward trunk neural crest (NC) cells. **(G-F)** qPCR expression analysis of representative lineage-specific, axial identity **(G)** and HOX genes **(F)** in NMP-derived trunk NC cultures treated with the indicated combinations of WNT-BMP agonists/antagonists.

We next examined the signalling pathway dependence of HOX gene expression during the transition of NMPs toward trunk NC cells, which involves a 5-day treatment with a lower amount of the WNT agonist CHIR and an intermediate level of BMP activity; the latter is achieved through the addition of a saturating amount of recombinant BMP4 together with the BMP type I receptor inhibitor DMH-1 to antagonistically modulate the effects of BMP4 (**Fig. 3E**) (Frith et al., 2018, Frith & Tsakiridis, 2019, Hackland et al., 2017). We examined the effect of perturbing the two main signalling pathways driving the specification of NC from NMPs (WNT and BMP) using agonist/antagonist combinations between days 3-8, in a manner similar to the day 0-3 treatments (**Fig. 3E**). Treatment of differentiating NMPs with XAV to inhibit WNT signalling in combination with BMP stimulation, resulted in a reduction in the expression of most HOX genes analysed albeit to a lesser degree compared to days 0-3 (**Fig. 3F,** dark blue vs purple bars). This was accompanied by a moderate increase in the expression of anterior (*OTX2*, *ETS1*) and decrease in posterior (*CDX2*) NC markers while the levels of neural/NC-specific transcripts appeared unaffected (**Fig. 3G,** dark blue vs purple bars). On the contrary, WNT stimulation in combination with the BMP inhibitor LDN193189 (LDN) had no effect or resulted in upregulation in the expression of most HOX genes relative to the WNT-BMP controls (**Fig. 3F,** light blue vs purple bars). Moreover, this treatment resulted in complete extinction of the expression of the definitive neural crest marker *SOX10* and a concomitant increase in the expression of the neural progenitor marker *SOX1* (**Fig. 3G,** light blue vs purple bars) pointing to a role for BMP signalling in steering NMPs/dorsal pre-neural progenitors toward a NC fate in agreement with previous observations (Leung et al., 2016). Taken together, our results demonstrate that HOX gene activation in hPSC-derived NMPs and its tight coupling to the early programming of a posterior axial identity in NMP-derived NC cells are primarily driven by a WNT-TBXT loop. However, the early dependence of HOX gene expression on WNT signalling appears to diminish with time as NMPs gradually differentiate toward NC.

We went on to examine whether the reliance of early trunk NC progenitor patterning on WNT signalling occurs *in vivo*. Homotopic grafting-based fate mapping has indicated that NM-potent cells in the lateral-most caudal epiblast of somitogenesis-stage mouse embryos (marked as “LE” in **Fig. 4**) can give rise to posterior NC (Wymeersch et al., 2016). We examined *R26-WntVis* embryos, in which graded nuclear EGFP expression relates to the degree of WNT signalling strength (Takemoto et al., 2016), at the time of posterior NC formation (Embryonic day (E) 8.75-E9.0, Theiler stage 13-14). In agreement with previous work (Ferrer-Vaquer et al., 2010), the highest levels of Wnt activity were confined within the posterior growth region, whereas anteriorly only cells in the head mesenchyme exhibited medium levels of EGFP (**Fig. 4A**). As the axis grows, posterior trunk tissue (such as the somites) were also found to display high EGFP expression (**Fig. 4A**). Wholemount immunostaining and 3D modelling of the highest EGFP-positive fraction (a-GFP^high^) demonstrated that NC-fated regions within the T(Brachyury)^+^ lateral-most caudal epiblast domain are marked by higher WNT signalling levels compared to more medial positions (dashed lines in **Fig. 4B**, **EV2A-B**). Expression of Tfap2a, a marker indicative of early NC specification (Mitchell et al., 1991, Rothstein & Simoes-Costa, 2020) was first detected within the Wnt^high^ LE domain at E9.0, whereas before it was only present in the non-neural ectoderm (compare arrows in **Fig. 4Ce-f** vs **4Cm-o**). At E8.75, Wnt activity levels appeared lower within committed, Sox9^+^ pre-migratory NC cells located in the dorsolateral neural tube (**Fig. 4Cc-d; EV2C**). However, E9.0 delaminating trunk NC and cranial NC cells were found to exhibit progressively elevated Wnt activity (**Fig. 4Cg-i** and **4CJ-l; EV2C**) reflecting the well-established role of this signalling pathway in promoting NC delamination and subsequent migration (Azambuja & Simoes-Costa, 2021, Burstyn-Cohen et al., 2004, Gandhi et al., 2020, Liu et al., 2013). Together these findings demonstrate that early specification of NM-potent TBXT^+^ trunk NC progenitors correlates with high WNT activity but this association becomes less prevalent as they gradually commit to a definitive, pre-migratory NC fate in line with our *in vitro* observations showing the early but progressively decreasing WNT-dependence of TBXT-driven posterior axial identity programming in NC cells.

**Figure 4.**
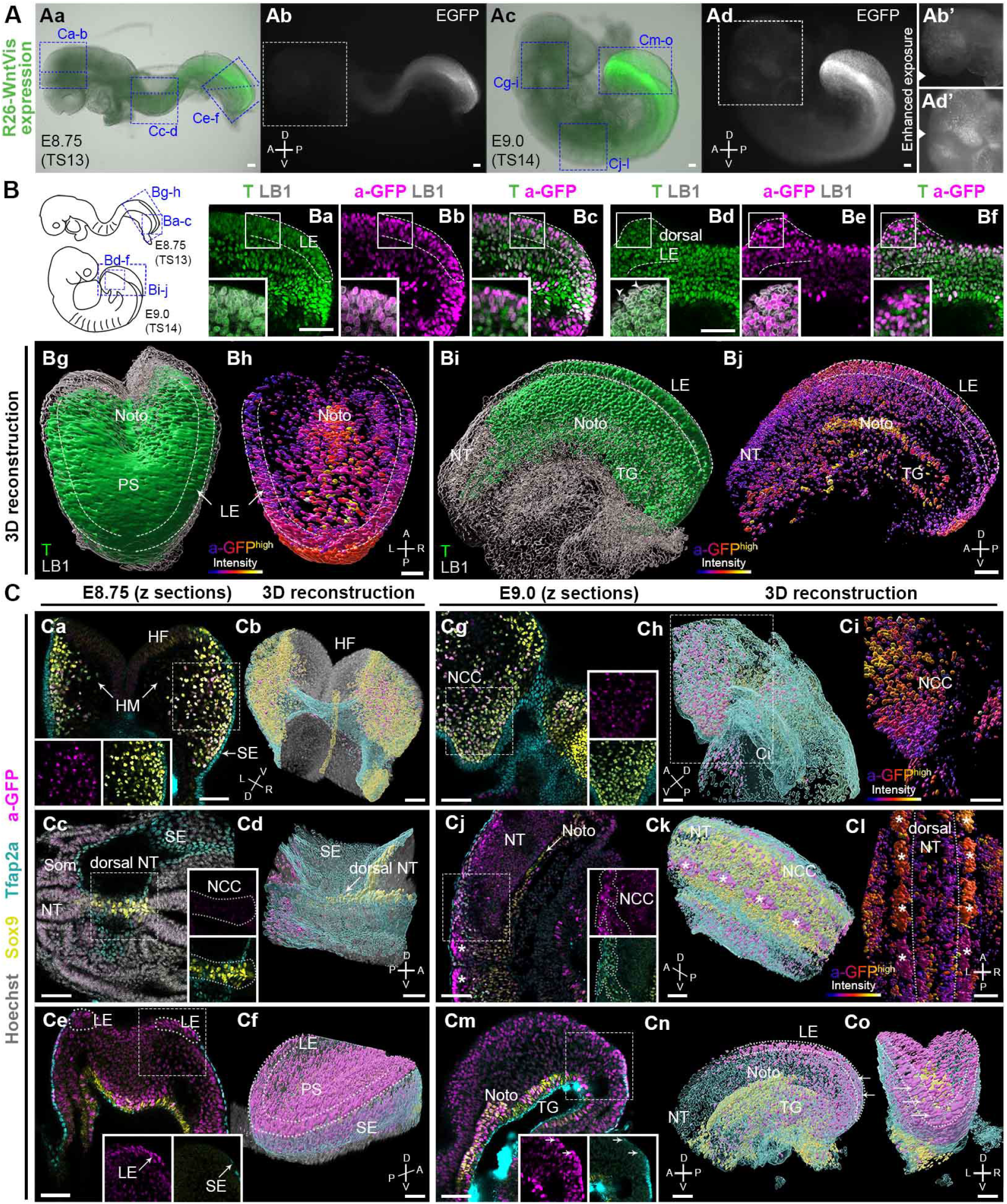
Wnt signalling dynamics during posterior neural crest emergence. **(A)** Fluorescence microscopy analysis of R26-WntVis mouse embryos at E8.75 (Theiler Stage; TS13) and E9.0 (TS14) showing graded responsiveness to Wnt signalling in the posterior growth region. **(B)** Sagittal sections of immunostained *R26-WntVis* tail buds showing a-GFP signal in the T^+^ lateral-most caudal epiblast (LE; dashed lines) at E8.75 (**Ba-c**) and E9.0 (**Bd-f**). Arrowheads indicate T^-^ surface ectoderm cells. (**Bg-j**) 3D reconstruction and processing of imaging data showing T^+^ volumes overlapping with a-GFP^high^ volumes (i.e. Wnt^high^ T^+^ cells) in the LE: posterior view at E8.75 (**Bg-h**) and lateral view at E9.0 (**Bi-j**). A cut-off was made to show only higher expressing a-GFP^+^ volumes (see **EV2**). Nuclei were defined by anti-Laminin B1 staining (LB1). **(C)** Wholemount immunostaining and corresponding 3D analysis showing NC derivatives and their progenitors at different sections of the rostrocaudal axis (regions corresponding to blue boxes in A). Insets corresponding to the dash line-boxed areas show the a-GFP and overlaid Sox9 and Tfap2a channels, respectively. (**Ca-b, Cg-i**) Sox9^+^ Tfap2a^+^ cells in the developing head mesenchyme. Sox9^+^ Tfap2a^+^ NC cells emerging from the E8.75 dorsal neural tube show low Wnt activity (**Cc-d**), whereas those at E9.0 generally exhibit a higher GFP signal (compare Cc to Cj). Asterisks mark dorsal somite cells. At E8.75, no Tfap2a^+^GFP^+^ cells were observed in the LE layer (**Ce-f**), whereas at E9.0 the caudalmost epiblast contains a population of Tfap2a^+^ GFP^+^ cells (arrows in Cm-o). HF, headfolds; HM, head mesenchyme; NCC, neural crest cells; Noto, notochord; NT, neural tube; PS, primitive streak; Som, somite; SE, surface ectoderm; TG, tail gut; A, anterior; P, posterior; D, dorsal; V, ventral; L, left; R, right. Scale bars=50 μm.

### TBXT controls posterior axial identity acquisition by influencing chromatin accessibility

We next examined whether TBXT controls HOX gene activation and potentiates the action of posteriorising extrinsic signals in CHIR-FGF-treated hPSCs via direct genomic binding or, indirectly, by regulating the expression of other key posteriorising regulators such as CDX2. To this end, we carried out chromatin immunoprecipitation followed by high-throughput sequencing (ChIP-seq) on hPSC-derived NMPs and undifferentiated hPSCs (control) to map TBXT targets genome-wide. We identified 24,704 TBXT-binding regions in NMPs (**Appendix Table S3, Fig. EV3A**), a large fraction of which were located within introns (~50%) and distal intergenic (~38%) regions (**Fig. EV3B**), reflecting similar findings on mouse T binding (Beisaw et al., 2018, Tosic et al., 2019). A small number (n=2088) of non-overlapping TBXT binding sites was also detected in undifferentiated hESCs, in line with the reported low-level expression of this transcription factor in mesoderm-biased pluripotent cells (Stavish et al., 2020, Tsakiridis et al., 2014). GO biological processes enrichment analysis of target genes associated with TXBT binding sites around the transcriptional start site (−2000 to +500 bp; 3084 peaks, 2841 genes) revealed an over-representation of established developmental regulators of A-P regionalisation (Benjamini–Hochberg P adj<0.05; **Fig. EV3C, Appendix Table S4**). Transcription factor motif enrichment analysis revealed that the TBXT binding regions in hPSC-derived NMPs were enriched, as expected, for the Brachyury consensus binding motif as well as other T-BOX binding motifs such as EOMES and TBX6 (**Fig. 5A, Appendix Table S5**), in agreement with previous findings in the mouse demonstrating that a fraction of the genomic targets of these transcription factors is also occupied by TBXT (Koch et al., 2017, Tosic et al., 2019). Other over-represented motifs included WNT signalling effectors (LEF1/TCF3) and Homeobox genes such as CDX factors (CDX2/4) and various HOX family members (**Fig. 5A, Appendix Table S5**) reflecting possible cooperative binding between these factors and TBXT in human NMPs. Comparison of TBXT ChiP-seq targets with our list of differentially expressed genes following Tet-induced TBXT knockdown in day 3 FGF-CHIR treated cultures (**Fig. 1**) revealed that the majority (~85%) of the downregulated (P adj <0.05, log2FC > |1|) genes, including HOX PG(1-9) members, pro-mesodermal factors and WNT-FGF signalling components, were directly bound by TBXT, while most (~60%) of their upregulated counterparts were not (**Fig. 5B, 5C, 5F, EV3D, Appendix Table S6)**. This indicates that TBXT acts primarily as a transcriptional activator of key downstream NMP/mesoderm regulators and HOX genes as previously reported (Amin et al., 2016, Beisaw et al., 2018, Guibentif et al., 2021, Koch et al., 2017, Lolas et al., 2014).

**Figure 5.**
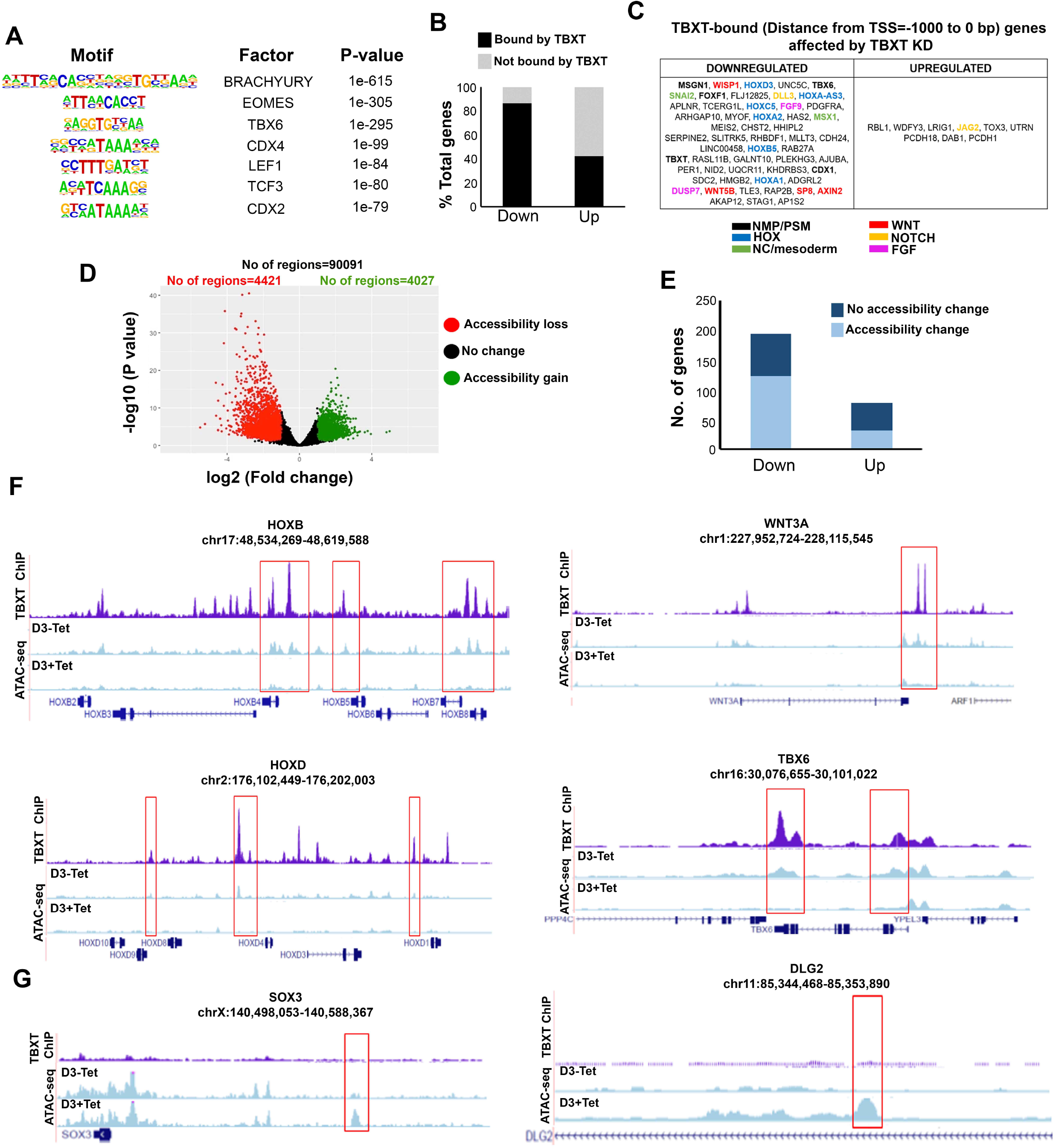
TBXT controls the HOX clock and WNT signalling in NMPs by influencing chromatin accessibility. **(A)** Representative transcription factor-binding motifs enriched in TBXT binding sites. **(B)** Graph showing the percentages of differentially expressed genes (Padj<0.05, log2FC> |1|) following TBXT knockdown during the transition of hESCs toward NMPs that are bound directly by TBXT. **(C)** Table showing all differentially expressed genes following TBXT knockdown that exhibit TBXT binding within their promoter region. **(D)** Volcano plot of differentially accessible ATAC seq peaks between TBXT-depleted and control NMPs. **(E)** Graph showing the number of significantly up- and down-regulated direct TBXT targets in relation to changes in chromatin accessibility associated with TBXT knockdown. **(F)** Correspondence between TBXT binding (ChiP) and chromatin accessibility changes (ATAC-seq) in the presence and absence of Tet in indicated HOX clusters, WNT/presomitic mesoderm-linked loci. **(G)** Correspondence between TBXT binding and chromatin accessibility changes in the presence and absence of Tet in indicated neural differentiation-linked loci.

To further dissect the impact of TBXT binding, we interrogated the chromatin accessibility landscape in TBXT-depleted and control hPSC-derived NMPs using ATAC-seq (assay for transposase accessible chromatin with high-throughput sequencing) (Buenrostro et al., 2013). We detected 4421 and 4027 unique regions corresponding to loss or gain of chromatin accessibility following Tet treatment, respectively (distance from TSS ranging from −1 to 1 kb, log2FC cutoff = 2 and p-value < 0.05) (**Fig. 5D, Appendix Table S7**). Genes associated with loss of chromatin accessibility were predominantly linked to A-P patterning, mesoderm specification and WNT signalling (**Appendix Table S7**). Moreover, the Tet-induced downregulation of most transcripts found to be TBXT targets (**Appendix Table S3**) was also accompanied by a significant loss of ‘open’ chromatin regions within the TBXT binding sites (**Fig. 5E, 5F, Appendix Table S7**). These included sites that were dispersed within all HOX clusters as well as key presomitic mesoderm regulators (e.g. TBX6, MSGN1) and WNT signalling components (e.g. WNT3A/8A, LEF1) (**Fig. 5F, EV3D**). On the contrary, there was no obvious correlation between upregulation of expression following TBXT knockdown and chromatin accessibility change (**Fig. 5E, Appendix Table S7**). Gain of accessible regions in TBXT-depleted hPSC-derived NMPs appeared to occur predominantly within/around loci linked to neural development (e.g. SOX3 and DLG2 (Tao et al., 2003, Wood & Episkopou, 1999)), most of which were not bound directly by TBXT (**Fig. 5G, Appendix Table S7**). Moreover, examination of transcription factor binding motifs (annotated using HOMER (Heinz et al., 2010); Benjamini–Hochberg P adj<0.05) showed that ATAC-seq sites marked by loss of chromatin accessibility following TBXT knockdown were uniquely enriched in CDX, T-box factor and HOX binding-associated DNA sequence elements whereas the regions marked by gain of chromatin accessibility were mainly characterised by motifs indicating binding of SOX, OCT/POU and Forkhead family transcription factors (**Fig. EV3E, Appendix Table S8**). Collectively, these data suggest that TBXT actively reconfigures the chromatin landscape in HOX genes and other axis elongation regulators, including WNT signalling components, by direct binding to key regulatory elements to promote the acquisition of a posterior axial character/mesoderm identity during the transition of pluripotent cells toward a caudal epiblast/NMP state.

### Posterior axial identity acquisition by NMP-derived pre-neural spinal cord cells is TBXT-independent and FGF-dependent

We next sought to examine whether early TBXT knockdown-triggered disruption of HOX gene expression/NMP induction also affects the CNS derivatives of NMPs. We thus generated early pre-neural spinal cord progenitors, which we have recently shown to give rise to posterior motor neurons of a thoracic axial identity, from TBXT shRNA hESCs (Wind et al., 2021). Cells were treated with CHIR-FGF2 for 3 days followed by their re-plating and further culture in the presence of CHIR and high FGF levels for 4 days combined with continuous Tet treatment to mediate TBXT knockdown (**Fig. 6A**). The levels of most HOX transcripts examined were largely unaffected in day 7 pre-neural spinal cord progenitors generated from TBXT-depleted cells, and similar to untreated controls (**Fig. 6B**). No dramatic changes in the expression of *CDX2* and *SOX2* (markers of an early spinal cord character at this stage) were observed (**Fig. 6C**). In line with our previous observations and published *in vivo* data (Gofflot et al., 1997, Nordstrom et al., 2006, Wind et al., 2021), we also detected low levels of TBXT transcripts in control cultures and, surprisingly, these were comparable to their Tet-treated counterparts (**Fig. 6C**). This finding suggests that selection of cells evading shRNA knockdown and maintaining low levels of *TBXT* may occur upon culture in pre-neural spinal cord cell-promoting culture conditions. Later addition of Tet at day 3, at the start of NMP differentiation toward pre-neural spinal cord cells, appeared to restore efficient TBXT knockdown (**Fig. 6D, F**) but, again, had no major impact on either HOX PG(1-9) gene expression or the levels of the early spinal cord/pre-neural transcripts *CDX2* and *NKX1-2* in the resulting day 7 cultures (**Fig. 6E, F**). These results indicate that, in contrast to NMP-derived NC, HOX expression dynamics/posterior axial identity in NMP-derived pre-neural spinal cord cells are unlikely to be controlled by TBXT.

**Figure 6.**
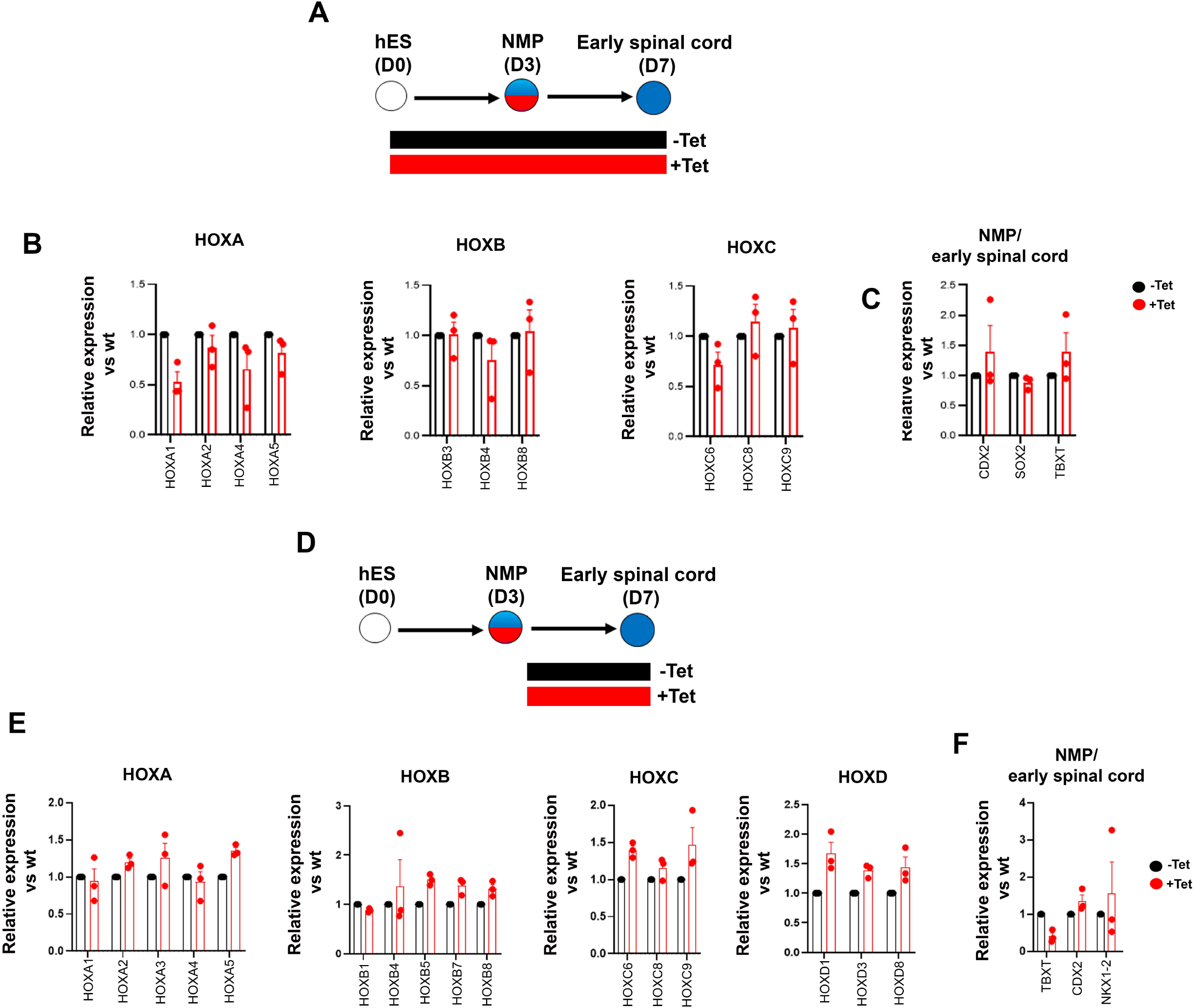
Posterior axial identity acquisition by NMP-derived pre-neural spinal cord cells is TBXT-independent. **(A, D)** Differentiation/treatment schemes associated with different time windows of TBXT knockdown during spinal cord differentiation from NMPs. **(B-C, E-F)** qPCR expression analysis of indicated HOX genes (**B, E**) and representative NMP/early spinal cord markers (**C, F**) in control vs Tet-treated NMP-derived early spinal cord progenitors corresponding to the Tet treatment regimens shown in A and D respectively.

Both WNT and FGF signalling have been previously implicated in the control of the Hox clock/early posterior patterning of spinal cord cells, through cooperation with the key axis elongation factor Cdx2 (Bel-Vialar et al., 2002, Lippmann et al., 2015, Liu et al., 2001, Mazzoni et al., 2013, Metzis et al., 2018, Mouilleau et al., 2021, Nordstrom et al., 2006, Olivera-Martinez et al., 2014, Takemoto et al., 2006, van de Ven et al., 2011). Our data so far have demonstrated an early requirement for WNT signalling in inducing a posterior axial identity/Hox activation during the transition of hPSCs toward NMPs. This reliance on WNT signalling diminishes during the transition of NMPs toward an NC fate. We tested whether this is also the case during the differentiation of NMPs toward pre-neural spinal cord progenitors (following their re-plating in the presence of CHIR-high FGF2 levels). We carried out signalling agonist/inhibitor combination experiments as described earlier (**Fig. 3**) treating differentiating cells either with the WNT inhibitor XAV in the presence of FGF2 or the FGF/MEK inhibitor PD03 in combination with CHIR between days 3-7 of differentiation (**Fig. 7A**). We found that WNT signalling inhibition in the presence of FGF activity during this time window does not appear to have a major effect on HOX gene expression or *CDX2* levels (blue bars in **Fig. 7B-D**). In contrast, blocking FGF signalling in combination with CHIR treatment resulted in a dramatic reduction in the expression of most HOX genes examined, particularly those belonging to PG(4-9), as well as *CDX2* (red bars, **Fig. 7B-D**). The NMP/pre-neural spinal cord marker *NKX1-2* was equally affected by the two treatments while the levels of later spinal cord markers (*PAX6*, *SOX1*) were modestly elevated (**Fig. 7C**). Collectively, these data suggest that HOX gene as well as CDX2 expression levels during the specification of early spinal cord cells from hPSC-derived NMPs rely primarily on FGF signalling and do not appear to depend on the action of an early NMP-based WNT-TBXT regulatory loop.

**Figure 7.**
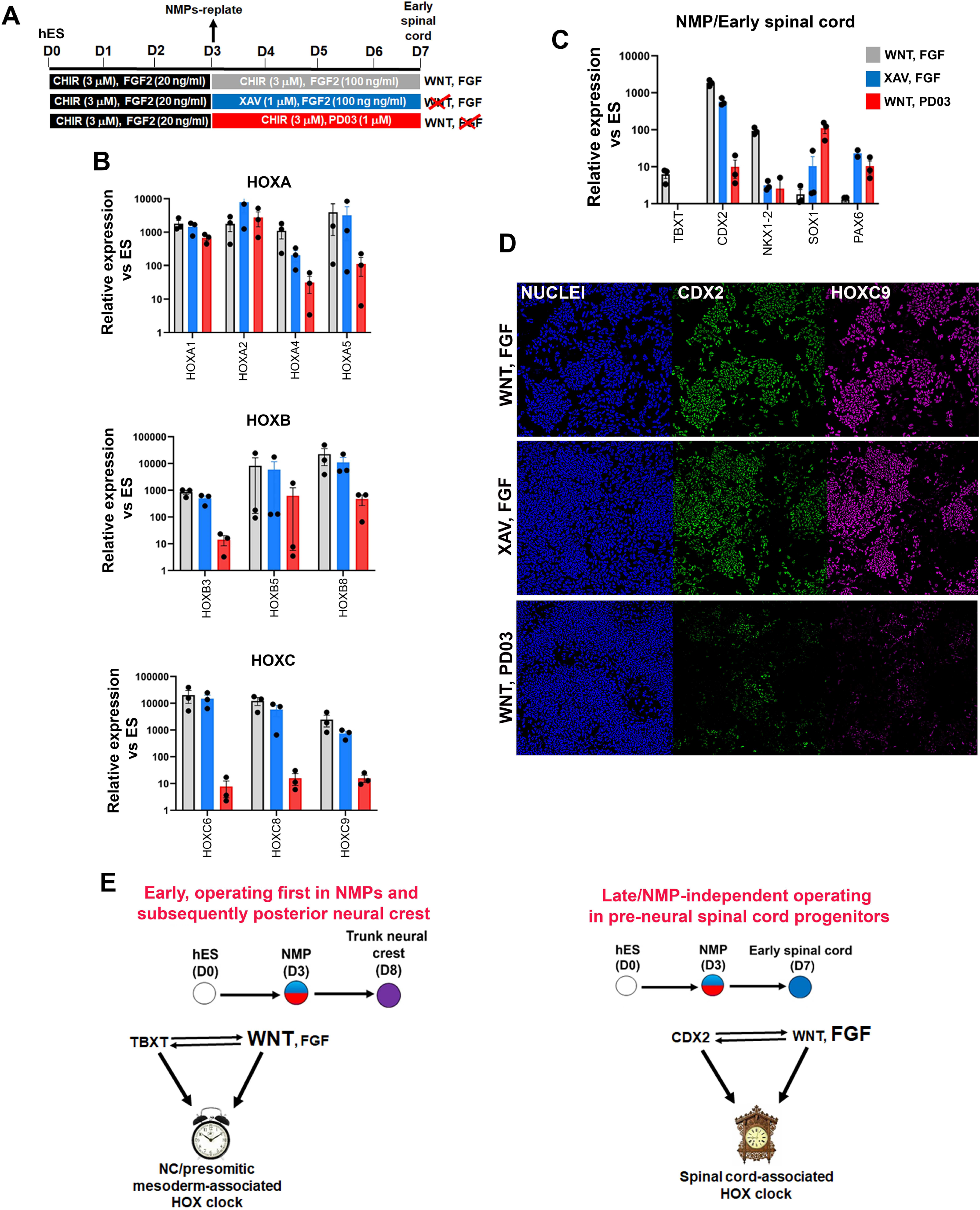
Posterior axial identity acquisition by NMP-derived spinal cord progenitors is FGF-dependent. **(A)** Scheme of treatments during the differentiation of hPSC-derived NMPs toward early spinal cord progenitors. **(B-C)** qPCR expression analysis of indicated HOX genes **(B)** and representative NMP/early spinal cord/neural markers **(C)** in NMP-derived early spinal cord cultures treated with the indicated combinations of WNT-FGF agonists/antagonists as depicted in A. **(D)** Immunofluorescence analysis of CDX2 and HOXC9 protein expression in NMP-derived early spinal cord cultures treated with the indicated combinations of WNT-FGF agonists/antagonists. **(E)** Proposed model for the transcriptional and signalling control of posterior axial identity/Hox clock in hPSC-derived NMPs and their derivatives.

## Discussion

A number of recent studies have demonstrated that the emergence of the trunk NC is tightly coupled to the induction of NM-potent progenitors localised in the post-gastrulation/somitogenesis stage posterior growth region of vertebrate embryos. Here we provide evidence indicating that the encoding of a posterior axial identity in human NC cells, embodied by the sequential activation of Hox genes and the induction of posterior markers, relies on their transition through this developmentally-plastic entity. This “primary regionalisation” process (Metzis et al., 2018) is mediated by the action of the pro-mesodermal transcription factor TBXT, which directs the operation of the Hox clock and the expression of posterior/presomitic mesoderm regulators together with WNT signalling effectors and, possibly, CDX2 (**Fig. 7E**). The early WNT dependence of trunk NC-associated Hox gene expression diminishes after NMP specification as definitive pre-migratory NC cells begin to emerge, before this signalling pathway becomes critical again in controlling later NC delamination/migration events. Strikingly, the expression of HOX genes, particularly those belonging to the central/posterior PG(4-9), in prospective posterior CNS spinal cord cells is mainly controlled by FGF signalling and appears unaffected by early HOX expression disruption events induced by TBXT depletion. This suggests that a separate NMP/TBXT/WNT-independent mechanism of Hox gene transcriptional control operates at a later post-NMP specification but pre-neural commitment stage, within this lineage (**Fig. 7E**).

Our proposed model synthesises previous findings showing that WNT-driven posterior axial identity acquisition in neural derivatives of NMPs takes place prior to neural induction (Lippmann et al., 2015, Metzis et al., 2018, Neijts et al., 2017, Neijts et al., 2016, Takemoto et al., 2006) with data illustrating that temporally discrete modes of trunk axial patterning in CNS spinal cord/NC rely on both FGF and WNT activities (Bel-Vialar et al., 2002, Delfino-Machin et al., 2005, Dunty et al., 2014, Gomez, 2019, Hackland et al., 2019, Liu et al., 2001, Mazzoni et al., 2013, Mouilleau et al., 2021, Muhr et al., 1999, Nordstrom et al., 2006, Sanchez-Ferras et al., 2016, Sanchez-Ferras et al., 2014, Sanchez-Ferras et al., 2012, van Rooijen et al., 2012, Zhao et al., 2014). Our work also reflects results showing the existence of distinct phases of Hox gene expression program implementation *in vivo*: (1) an early “plastic” phase linked to A-P patterning of multipotent axial progenitors/NMPs within their posterior niche and (2) a later phase, which marks the instalment and fixing of lineage-specific Hox gene expression patterns and the sharpening of their final boundaries in the neural and mesodermal derivatives of axial progenitors as and after they exit the posterior growth zone (Ahn et al., 2014, Brend et al., 2003, Charite et al., 1998, Deschamps & Wijgerde, 1993, Forlani et al., 2003, Hayward et al., 2015, McGrew et al., 2008, Wymeersch et al., 2019).

We demonstrate that TBXT function is crucial for proper collinear activation of Hox genes during NMP induction from hPSCs *in vitro* by channelling extrinsic WNT activity toward the establishment of a “posteriorising”/pro-mesodermal niche (in line with previous data (Martin & Kimelman, 2010)) that is essential for the subsequent transduction of a posterior character to NMP-derived NC cells. This prospective TBXT-WNT-driven transmission of positional information via an early axial progenitor intermediate and the potential inductive interaction between nascent presomitic mesoderm and NC progenitors reinforces previous observations on the effect of non-organiser mesoderm and vertical signalling as determinants of HOX gene expression and axial identity in early neural cells (Bardine et al., 2014, Forlani et al., 2003, Grapin-Botton et al., 1997). However, further work is required to disentangle the cell vs non-cell autonomous effects of TBXT on the A-P patterning of NC cells.

Our results are in line with previous studies reporting that TBXT attenuation in hPSC-derived mesoderm progenitors and spinal cord organoids as well as in Xenopus morphant and mouse *Tc* (Curtailed) mutant embryos abolishes Hox gene expression (Faial et al., 2015, Libby et al., 2021, Lolas et al., 2014, Wacker et al., 2004). Moreover, they further expand the repertoire of actions that orchestrate embryonic axis elongation/axial progenitor ontogeny and are exerted via the regulatory axis centred on WNT signalling, TBXT and the Hox clock (Denans et al., 2015, Mariani et al., 2021, Ye et al., 2021, Ye & Kimelman, 2020). Interestingly, global Hox gene expression appears minimally affected in some Brachyury mutants (such as those with the *T^2J/2J^* genotype (Koch et al., 2017, Tosic et al., 2019)), as well as in *T*^-/-^:: Wild type chimeras (Guibentif et al., 2021) suggesting that the nature of the *T* gene mutation and non-cell autonomous rescue effects from the surrounding wild-type environment, similar to those described previously in the case of grafted *Cdx* mutant axial progenitors (Bialecka et al., 2010), are critical actors in influencing the effect of Brachyury on its target genes. Moreover, the developmental arrest exhibited by *T* mutant embryos around the time of trunk NC emergence and the associated lack of posterior axial tissue precludes detailed assessment of Hox gene expression in *T* null NC cells.

TBXT has been shown to regulate its downstream mesoderm/axis elongation-associated targets in mouse embryos by promoting chromatin remodelling and directing permissive chromatin modifications in key regulatory elements (Amin et al., 2016, Beisaw et al., 2018, Koch et al., 2017, Tosic et al., 2019). Our data confirm that this is also the case during the transition of hPSCs toward NMPs and reveal that TBXT additionally contributes to global control of Hox cluster transcription in a similar way and in line with previous reports showing direct TBXT binding in Hox loci during hPSC differentiation (Faial et al., 2015). We propose that concomitant activation of the Hox clock and induction of WNT signalling components, via TBXT-driven chromatin landscape reconfiguration, comprise a critical early step in primary A-P regionalisation and the transition of pluripotent cells toward a caudal epiblast/axial progenitor state. This is supported by our demonstration that TBXT knockdown results in the acquisition of an anterior epiblast/AVE identity associated with WNT antagonism/head formation-promoting activity as well as increased chromatin accessibility in regulatory elements controlling pro-neural differentiation genes. Previous data on the crucial role of Brachyury in counteracting the default neurectodermal differentiation of mouse pluripotent cells by altering chromatin accessibility in key enhancers also point to such a model (Tosic et al., 2019). Moreover, early A-P regionalisation driven by the TBXT-WNT-HOX axis is likely to involve the cooperative action of CDX factors and the participation of other key transcriptional factors such as EOMES and SOX2 given their previously reported functions in the mouse (Amin et al., 2016, Blassberg, 2020, Metzis et al., 2018, Neijts et al., 2017, Neijts et al., 2016, Tosic et al., 2019). Thus, additional thorough dissection of the individual roles of these factors via loss-/gain of-function approaches is required in order to elucidate their crosstalk as they fine-tune the balance between pluripotency exit, cell fate decision making and adoption of a posterior character in human cells.

In summary, we provide a mechanistic insight into the cellular and molecular basis of posterior axial identity acquisition during hPSC differentiation. Our data demonstrate a novel role for TBXT in controlling Hox gene expression and early posteriorisation supporting the idea that A-P patterning of at least some axial progenitor derivatives, such as the trunk neural crest, occurs prior to their specification, within their multipotent precursors. We speculate that the close links between TBXT-driven posterior axial identity programming in the neural crest and NMP ontogeny may explain some cases of spina bifida observed in individuals carrying mutations within the TBXT locus (Agopian et al., 2013, Carter et al., 2011, Fellous et al., 1982, Morrison et al., 1996, Shields et al., 2000), especially in light of the potential involvement of impaired NC specification and HOX gene dysregulation in neural tube defects (Anderson et al., 2016, Degenhardt et al., 2010, Poncet et al., 2020, Rochtus et al., 2015, Yu et al., 2019).

## Materials and Methods

### Cell culture and differentiation

Use of hPSCs has been approved by the Human Embryonic Stem Cell UK Steering Committee (SCSC15-23). The *TBXT* and *B2M* shRNA sOPTiKD hESC lines (H9 background) (Bertero et al., 2016) were employed for all TBXT loss-of-function and sequencing experiments whereas signalling agonist/antagonist treatments were performed in H9 hESCs and SFCi55-ZsGr human induced PSCs (Lopez-Yrigoyen et al., 2018, Thomson et al., 1998). All cell lines were cultured routinely in feeder-free conditions in either Essential 8 (Thermo Fisher or made in-house) or mTeSR1 (Stem Cell Technologies) medium on laminin 521 (Biolamina) or Geltrex LDEV-Free reduced growth factor basement membrane matrix (Thermo Fisher). Cells were passaged twice a week after reaching approximately 80% confluency using PBS/EDTA as a passaging reagent. TBTX inducible knockdown in the *TBXT* shRNA sOPTiKD hESC line was achieved using tetracycline hydrochloride (Merck Life Science) at 1μg/ml as described previously (Bertero et al., 2016). hESCs were cultured in the presence/absence of tetracycline for 2 days prior to the initiation of differentiation and the tetracycline treatment was continued throughout the differentiation for the periods indicated in the results section/schemes.

For NMP differentiation, hPSCs (70-80% confluent) were dissociated using Accutase solution (Merck Life Science) and plated at a density of approximately 50,000 cells/cm^2^ on vitronectin (Thermo Fisher)-coated culture plates in N2B27 basal medium containing 50:50 DMEM F12 (Merck Life Science / Neurobasal medium (Gibco) and 1x N2 supplement (Gibco), 1x B27 (Gibco), 1x GlutaMAX (Gibco), 1x MEM NEAA (Gibco), 2-Mercaptoethanol (50 μM, Gibco). N2B27 medium was supplemented with CHIR99021 (3 μM, Tocris), FGF2 (20 ng/ml, R&D Systems) and Rho-Associated Coil Kinase (ROCK) inhibitor Y-27632 2HCl (10 μM, Adooq Biosciences) with the latter being withdrawn from the differentiation medium after the first day of NMP induction. For TBXT inducible knockdown, NMP medium was supplemented with 1μg/ml tetracycline hydrochloride and replenished every other day. For early spinal cord progenitor differentiation, day 3 hPSC-derived NMPs were dissociated into single cell suspension using Accutase and re-plated at a density of 37,500 cells/cm^2^ on Geltrex-coated culture plates in N2B27 containing FGF2 (100 ng/ml), CHIR99021 (3 μM) and ROCK inhibitor Y-27632 2HCl (10 μM; for the first day only) in the presence or absence of tetracycline hydrochloride (1μg/ml). Medium was replaced every other day until day 7 of differentiation. For trunk neural crest differentiation, day 3 hPSC-derived NMPs were dissociated using Accutase and re-plated at a density of 50,000 cells/cm^2^ on Geltrex-coated culture plates directly into neural crest inducing medium consisting of DMEM/F12, 1x N2 supplement, 1x GlutaMAX, 1x MEM NEAA, the TGF-beta/Activin/Nodal inhibitor SB-431542 (2 μM, Tocris), CHIR99021 (1 μM), BMP4 (15ng/ml, Thermo Fisher), the BMP type-I receptor inhibitor DMH-1 (1 μM, Tocris) and ROCK inhibitor Y-27632 2HCl (10 μM). The medium was refreshed every two days without ROCK inhibitor and was supplemented with 1μg/ml tetracycline hydrochloride throughout the differentiation for tet-mediated inducible TBXT knockdown. Trunk neural crest cells were analysed at day 8 of differentiation. The following signalling pathway inhibitors were employed: the WNT antagonist tankyrase inhibitor XAV 939 (1 μM, Tocris), the MEK1/2 inhibitor PD0325901 (1 μM, Merck), LDN-193189 (100 nM, Tocris).

### Immunofluorescence and imaging

Cells were fixed in 4% PFA for 10 minutes at room temperature, rinsed twice with PBS and permeabilised with 0.5% Triton X-100 in PBS containing 10% Foetal Calf Serum (FCS) and 0.1% Bovine Serum Albumin (BSA) for 15 minutes. Blocking was then performed in blocking buffer consisting of 10% FCS, 0.1% BSA in PBS at room temperature for 1-2 hours or overnight at 4°C. Primary antibodies were diluted in blocking buffer and cells were incubated with primary antibodies overnight at 4oC. Following three washes, cells were incubated with secondary antibodies diluted in blocking buffer for 1-2 hours at room temperature and in the dark. Cell nuclei were counterstained with Hoechst (Thermo Fisher, 1:1000) and fluorescent images were acquired using the InCell Analyser 2200 system (GE Healthcare). Images then were processed in Fiji (Schindelin et al., 2012) or Photoshop (Adobe) using identical brightness/contrast settings to allow comparison between different treatments. At all times, the positive/negative threshold was set based using a sample incubated with secondary antibody only. The following primary antibodies were employed: anti-TBXT (Abcam, ab209665; 1:1000); anti-CDX2 (Abcam, ab76541; 1:200); anti-HOXC9 (Abcam, ab50839; 1:50); anti-SOX10 (Cell Signalling Technology, D5V9L; 1: 500).

For quantification of TBXT protein expression in NMP-like cells, nine random fields per experiment were scored (two biological replicates) and processed in the image analysis software CellProfiler (Carpenter et al., 2006). Nuclei were first identified using Hoechst staining and subsequently mapped onto the other fluorescent channels for single-cell fluorescence intensity quantification. Cells stained with secondary antibody only were used as negative control to set the negative/positive threshold. A histogram was created using GraphPad Prism (GraphPad Software). For quantification of SOX10 and HOXC9 protein expression in trunk neural crest, 5 random fields per experiment were scored (3 biological replicates) and processed as described above.

### Immunofluorescence analysis of wholemount embryos and 3D reconstruction

E8.75 and E9.0 (TS13-14) *R26-WntVis* embryos were dissected in-house prepared M2 medium (Nowotschin et al., 2010). Embryos with yolk sac and amnion removed were fixed for 25-30 minutes at 4°C in 4% PFA in PBS, followed by permeabilisation in 0.5% Triton X-100 in PBS for 15 minutes, then incubated in 0.5 M glycine in PBS in 0.1% Triton X-100 (PBST) for 20 minutes, after which they were washed three times in PBST and blocked overnight at 4°C in 10% serum (Merck) in PBS/0.3% TritonX100. Both primary and Alexa Fluor^®^-conjugated secondary antibodies (Thermo Fisher Scientific and Alexa Fluor^®^ 647 donkey-anti-chicken [#703-606-155] from Jackson Immunoresearch; all at 2 μg/ml final concentration) were diluted in blocking buffer and samples were incubated for 48 hours on a rocking platform at 4°C. A minimum of four 25-minute washes were performed with PBST on a rocking platform at room temperature after the primary and secondary antibody incubation. Antibodies used (supplier, final concentration): anti-GFP (Abcam; ab13970; 10 μg/ml); anti-Brachyury (R&D; AF2085; 1 μg/ml); anti-LB1 (Abcam; ab16048; 1:800-1:1000); anti-Sox9 (Merck, AB5535; 2 μg/ml); anti-Tfap2a (DSHB, 3B5, 3.1μg/ml). Embryos were counterstained with Hoechst33342 (Thermo Fischer Scientific; 5 μg/ml). Confocal microscopy was performed after dehydration through an increasing methanol/PBS series (25%, 50%, 75%, 10 min each) and two 5 min washes in 100% methanol and clearing in 1:1 v/v methanol/BA:BB (2:1 benzyl alcohol:benzyl benzoate; Sigma), and two washes in 100% BA:BB. Embryos were imaged in BA:BB under a LSM800 confocal system with Airyscan and GaAsP detectors (Zeiss). Two to three replicate embryos were imaged per staining and stage. Wholemount immunostaining data was processed using Zeiss or Imaris software (Oxford Instruments) using channel alignment, background subtraction and deconvolution tools. In Imaris, 3D volumes were created from single channels. These volumes represent positive stained areas, and - as volumes were allowed to merge - they do not represent cells. As many cells in the tail bud have some level of EGFP expression, only the fraction of a-GFP-stained cells with higher intensity (a-GFP^high^) were displayed in the analyses for clarity reasons (compare a-GFP signal to 3D volumes in **EV2B-D**). The filtering of a-GFP^low^ 3D volumes was kept constant between different regions of individual embryos.

### Quantitative real time PCR

Total RNA was extracted using the total RNA purification kit (Norgen Biotek) following manufacturer’s instructions. CDNA preparation was completed using the High-Capacity cDNA Reverse Transcription kit (Thermo Fisher). Quantitative real-time PCR was carried out using the QuantStudio 12 K Flex (Applied Biosystems) thermocycler in combination with the Roche UPL system and the TaqMan Fast Universal PCR Master Mix (Applied Biosystems). Primer sequences are shown in Appendix Table S9.

### Mouse husbandry

All mice were maintained on a 12 hr-light/12 hr-dark cycle and housed in 18–23 °C with 40–60% humidity. Homozygote *R26-WntVis* mice were obtained from the Laboratory for Animal Resources and Genetic Engineering at the RIKEN Center for Biosystems Dynamics Research, Kobe, Japan (accession no. CDB0303K; RBRC10162). Homozygote *R26-WntVis* males were crossed with ICR females (JAX stock #009122, The Jackson Laboratory); all experiments were performed on heterozygote embryos. For timed matings, noon on the day of finding a vaginal plug was designated E0.5. All animal experiments were approved by the Institutional Animal Experiments Committee of RIKEN Kobe Branch. Mice were handled in accordance with the ethics guidelines of the institute

### RNA-seq

#### Sample preparation

Total RNA was harvested from day 3 NMPs obtained from *TBXT* shRNA sOPTiKD hESC in the presence and absence of Tetracycline (three biological replicates) using the total RNA purification plus kit (Norgen BioTek) according to the manufacturer’s instructions. Sample quality control, library preparation and sequencing were carried out by Novogene (http://en.novogene.com). Library construction was carried out using the NEB Next Ultra RNA Library Prep Kit and sequencing was performed using the Illumina NoveSeq platform (PE150). Raw reads were processed through FastQC v0.11.2 (https://www.bioinformatics.babraham.ac.uk/projects/fastqc/) and Trim Galore (https://www.bioinformatics.babraham.ac.uk/projects/trim_galore/). Reads were aligned using STAR v2.4.2a (Dobin et al., 2013) to the human reference genome assembly GRCh38 (Ensembl Build 79) in the two-pass mode. RSEM v1.3.0 (Li & Dewey, 2011) was used to extract expected gene counts, where genes expressed in < 3 samples or with total counts ≤ 5 among all samples were excluded. We identified genes showing significant differential expression with DESeq2 (Love et al., 2014), with log2FoldChange> |0.5| and Benjamini–Hochberg-adjusted P<0.05. Data were deposited to GEO (Accession number: GSE184622).

### Chip-seq

Chromatin immunoprecipitation followed by next-generation sequencing (ChIP-seq) was performed on approximately 10 million cells/sample (three pooled replicates per sample), fixed with 1% formaldehyde solution (11% formaldehyde, 0.1M NaCl, 1mM EDTA (pH 8.0), 50mM HEPES (pH 7.9)) for 15 minutes at room temperature on a shaking apparatus. Fixation was quenched with 125mM of glycine (1/20 volume of 2.5M stock) for 5 minutes and then adherent cells were scraped thoroughly from the culture surface. Cells were washed, centrifuged at 800 x g for 10 minutes at 4°C and pellets were resuspended in 10ml chilled PBS-Igepal (1X PBS (pH 7.4), 0.5% Igepal CA-630). This pellet wash was repeated and cells were resuspended in 10 ml chilled PBS-Igepal and 1mM PMSF. Samples were centrifuged at 800 x g for 10 minutes at 4°C for a third time, after which the supernatant was removed and pellets were snap-frozen on dry ice and stored at −80°C. Samples were sent to Active Motif (Carlsbad, CA) for ChIP-seq. Active Motif (https://www.activemotif.com) prepared the chromatin, performed ChIP reactions, generated libraries, sequenced them and performed basic data analysis. Chromatin was isolated by adding lysis buffer, followed by disruption with a Dounce homogenizer. Lysates were sonicated and the DNA sheared to an average length of 300-500 bp with Active Motif’s EpiShear probe sonicator (cat# 53051). Genomic DNA (Input) was prepared by treating aliquots of chromatin with RNase, proteinase K and heat for de-crosslinking, followed by SPRI beads clean up (Beckman Coulter) and quantitation by Clariostar (BMG Labtech). Extrapolation to the original chromatin volume allowed determination of the total chromatin yield. An aliquot of chromatin (50 μg) was precleared with protein G agarose beads (Invitrogen) and genomic DNA regions of interest were isolated using 4 μg of antibody against Brachyury (R&D Systems, cat# AF2085, lot# KQP0719121). Complexes were washed, eluted from the beads with SDS buffer, and subjected to RNase and proteinase K treatment. Crosslinks were reversed by incubation overnight at 65°C, and ChIP DNA was purified by phenol-chloroform extraction and ethanol precipitation. Quantitative PCR (QPCR) reactions were carried out in triplicate on specific genomic regions using SYBR Green Supermix (Bio-Rad). The resulting signals were normalized for primer efficiency by carrying out QPCR for each primer pair using Input DNA. Illumina sequencing libraries were prepared from the ChIP and Input DNAs by the standard consecutive enzymatic steps of end-polishing, dA-addition, and adaptor ligation. Steps were performed on an automated system (Apollo 342, Wafergen Biosystems/Takara). After a final PCR amplification step, the resulting DNA libraries were quantified and sequenced on Illumina’s NextSeq 500 (75 nt reads, single end). Reads were aligned to the human genome (hg38) using the BWA algorithm (default settings) (Li & Durbin, 2009). Duplicate reads were removed and only uniquely mapped reads (mapping quality >25) were used for further analysis. Alignments were extended in silico at their 3’-ends to a length of 200 bp, which is the average genomic fragment length in the size-selected library, and assigned to 32-nt bins along the genome. The resulting histograms (genomic “signal maps”) were stored in bigWig files. Peak locations were determined using the MACS algorithm (v2.1.0) (Zhang et al., 2008) with a cutoff of p-value = 1e-7. Peaks that were on the ENCODE blacklist of known false ChIP-Seq peaks were removed. Signal maps and peak locations were used as input data to Active Motifs proprietary analysis program, which creates Excel tables containing detailed information on sample comparison, peak metrics, peak locations and gene annotations. Motif analysis was carried out using the Homer software (Heinz et al., 2010). Regions of 200 bp surrounding the summit of the top 2,500 peaks (based on MACS2 p-values) were analysed. Data were deposited to GEO (Accession number: GSE184622).

### ATAC-seq

Day 3 NMPs (50,000 cells) obtained from *TBXT* shRNA sOPTiKD hESC in the presence and absence of Tetracycline (three biological replicates) were harvested and samples were prepared using the Illumina Tagment DNA Enzyme and Buffer Small Kit (Illumina), 1% Digitonin (Promega) and EDTA-free Protease Inhibitor cocktail (Roche). Following DNA purification with the MinElute kit eluting in 12 μl, 1 μl of eluted DNA was used in a quantitative PCR (qPCR) reaction to estimate the optimal number of amplification cycles. The remaining 11 μl of each library were amplified for the number of cycles corresponding to the Cq value (i.e., the cycle number at which fluorescence has increased above background levels) from the qPCR. Library amplification was followed by AMPure beads (Beckman Coulter) size selection to exclude fragments smaller than 150bp and larger than 1,200 bp. Library amplification was performed using custom Nextera primers. DNA concentration was measured with a Qubit fluorometer (Life Technologies) and library profile was checked with Bioanalyzer High Sensitivity assay (Agilent Technologies). Libraries were sequenced by the Biomedical Sequencing Facility at CeMM (Research Center for Molecular Medicine of the Austrian Academy of Sciences, Vienna) using the Illumina HiSeq 3000/4000 platform and the 50-bp single-end configuration. Base calling was performed by Illumina Real Time Analysis (v2.7.7) software and the base calls were converted to short reads using the IlluminaBasecallsToSam tool from the Picard toolkit (v2.19.2) (“Picard Toolkit.” 2019. Broad Institute, GitHub Repository. http://broadinstitute.github.io/picard/; Broad Institute). Sequencing adapters were removed, and the low-quality reads were filtered using the fastp software (v 0.20.1) (Chen et al., 2018). Alignment of the short reads on GRCh38 was performed using Bowtie2 (v2.4.1) (Langmead & Salzberg, 2012) with the “-very-sensitive” parameter. PCR duplicates were marked using samblaster (v0.1.24) (Faust & Hall, 2014), and the reads mapped to the ENCODE black-listed (Amemiya et al., 2019) regions were discarded prior to peak calling. To detect the open chromatin regions, we performed peak calling using the MACS2 (v2.2.7.1) (Zhang et al., 2008) software with the “--nomodel”, “--keep-dup auto” and “--extsize 147” options. Peaks in the format of bed files were analysed for differential analysis to compare signals corresponding to the + vs –Tet samples using the GUAVA software. Differential peaks with a distance from TSS ranging from −1 to 1 kb, log2FC cutoff = 2 and p-value < 0.05 were extracted. Finally, HOMER findMotifs (v4.11) (Heinz et al., 2010)was used for motif enrichment analysis over the detected open chromatin regions. Data were deposited to GEO (Accession number: GSE184227).

## Acknowledgements

We wish to thank Prof. Ludovic Vallier (University of Cambridge) for providing the *TBXT* and *B2M* shRNA sOPTiKD hESC lines and Prof. Lesley Forrester (University of Edinburgh for providing the SFCi55-ZsGr iPSC line. We would also like to thank the Biomedical Sequencing Facility at CeMM/Michael Schuster and Leonardo Mottta, Estelle Suaud/Active Motif for assistance with the ATAC-seq and ChiP-seq experiments respectively. Finally, we are grateful to James Briscoe, Fay Cooper, Vicki Metzis, Matt Towers and Val Wilson for critical reading of the manuscript.

## Author contributions

Conceptualization: AT; Data curation: IG, IM, CB; Formal analysis: AG, CS, IG, IM; Funding acquisition: AT, MRG, MT, FJW; Investigation: AG, CS, IG, IM, FJW, MW, TJRF, MG, AB, AT; Methodology: AG, CS, IG, IM, FH, CB, AT; Project administration: AT; Resources: AB; Supervision: AT; Visualization: AG, CS, IG, IM, FJW, AT; Writing – original draft: AT; Writing – review & editing: AG, CS, IG, IM, FJW, MW, TJRF, AB, FH, CB, MRG, MT, AT.

## Tables and their legends

**Appendix Table S1.** Significantly up- and downregulated transcripts in Tet-treated, TBXT-depleted hESC-derived NMPs.

**Appendix Table S2.** List of GO terms and corresponding gene lists enriched in Tet-treated, TBXT depleted hESC-derived NMPs

**Appendix Table S3.** List of all genomic regions (Intervals) with peak p-value below the applied threshold bound by TBXT in hESC-derived NMPs and undifferentiated hESCs.

**Appendix Table S4.** List of GO terms and corresponding gene lists associated with TBXT binding sites in hPSC-derived NMPs.

**Appendix Table S5.** List of known HOMER database motifs enriched in TBXT binding sites in hESC-derived NMPs.

**Appendix Table S6.** List of TBXT target genes which are differentially expressed following Tet treatment (P adj <0.05 log2FC > |0.5|) and Gene Ontology Biological Processes enrichment analysis. Genes are listed in relation to the genomic position (in relation to TSS) of TBXT binding within their proximity. Blue highlight denotes downregulation while red represents upregulation in expression.

**Appendix Table S7.** List of ATAC-seq peaks associated with gain or loss of chromatin accessibility following TBXT depletion in hESC-derived NMPs. Gene Ontology Biological Processes enrichment analysis, list of HOX genes as well as other genes (P adj <0.05 log2FC > |1|) affected by TBXT depletion and are associated with changes in chromatin accessibility are also included.

**Appendix Table S8.** List of transcription factor DNA binding motifs enriched in ATAC-seq sites associated with chromatin accessibility gain, chromatin accessibility loss or both.

**Appendix Table S9.** List of primers used.

## Expanded View Figure legends

**Figure EV1. Effect of tetracycline treatment on control B2M shRNA hESC-derived NMPs. (A)** Immunofluorescence analysis of the expression of TBXT in control B2M shRNA hESC-derived NMPs in the presence and absence of tetracycline (Tet). **(B)** qPCR expression analysis of indicated HOX genes in control vs Tet-treated NMPs generated from B2M shRNA hESCs.

**Figure EV2. Wnt signalling dynamics during posterior neural crest emergence.**

Top: Schematics showing location and orientation of immunostaining data in E8.75 (TS13) and E9.0 (TS14) embryos. **(A)** Confocal sections of wholemount immunostaining shows graded responsiveness to Wnt signalling in T^+^ lateral-most caudal epiblast (LE, indicated by dashed lines). (**B-C**) Additional 3D views to *R26-WntVis* tail buds shown in **Fig.4B**: wholemount a-GFP staining (**Ba, Ca**), cut-off and mean intensity of GFP^high^ volumes (3D vol) (**Bb, Cb**) and overlap of Wnt^high^ and T^+^ volumes in the LE (dashed lines; **Bc, Cc**). The colour scale shows mean pixel intensity of a-GFP^high^ volumes, their numbers denoting the outermost values. **(D)** Comparison between a-GFP wholemount immunostaining data (magenta) to a-GFP^high^ 3D volume calculation (pink) and their mean pixel intensity (fire scale). (**Da-c)** Frontal view of the head region: a-GFP^low^ cells can be seen in the headfolds (HF; dotted lines) but were largely excluded from the analysis (arrowheads). **(Dd-f)** Dorsal view on the neural tube (NT, dotted lines). **(Dg-I, Dp-r)** Lateral view of the tail bud showing the LE (dotted lines). **(Dj-l)** Lateral view of the E9.0 head mesenchyme shows a-GFP^low^ cells are present in the hindbrain (HB, dotted lines). Note the gradient in a-GFP^+^ in neural crest cells (NCC; **Dl**). **(Dm-o)** E9.0 posterior axis shows high reporter expression in dorsal somites, graded from anterior to posterior (asterisks). Noto, notochord; PS, primitive streak; PSM, presomitic mesoderm; Som, somite; SE, surface ectoderm; TG, tail gut; A, anterior; P, posterior; D, dorsal; V, ventral; L, left; R, right. Scale bars=50 μm.

**Figure EV3. Effect of TBXT binding on chromatin accessibility. (A)** Average density plot of tag distributions across peak regions corresponding to the NMP, hES and input samples. **(B)** Genomic distribution of TBXT-bound sites in hPSC-derived NMPs. **(C)** Gene ontology biological processes enrichment analysis of target genes associated with TXBT binding sites around their transcriptional start site (−2000 to +500 bp). **(D)** Correspondence between TBXT binding (ChiP) and chromatin accessibility changes (ATAC-seq) in the presence and absence of Tet in indicated HOX and WNT-linked loci. **(E)** Venn diagram showing the overalp between transcription factor DNA binding motifs in genomic regions associated with chromatin accessibility gain and loss following TBXT knockdown.

