## Supplementary figures and images for "Early anteroposterior regionalisation of human neural crest is shaped by a pro-mesodermal factor"

### Fig. EV1

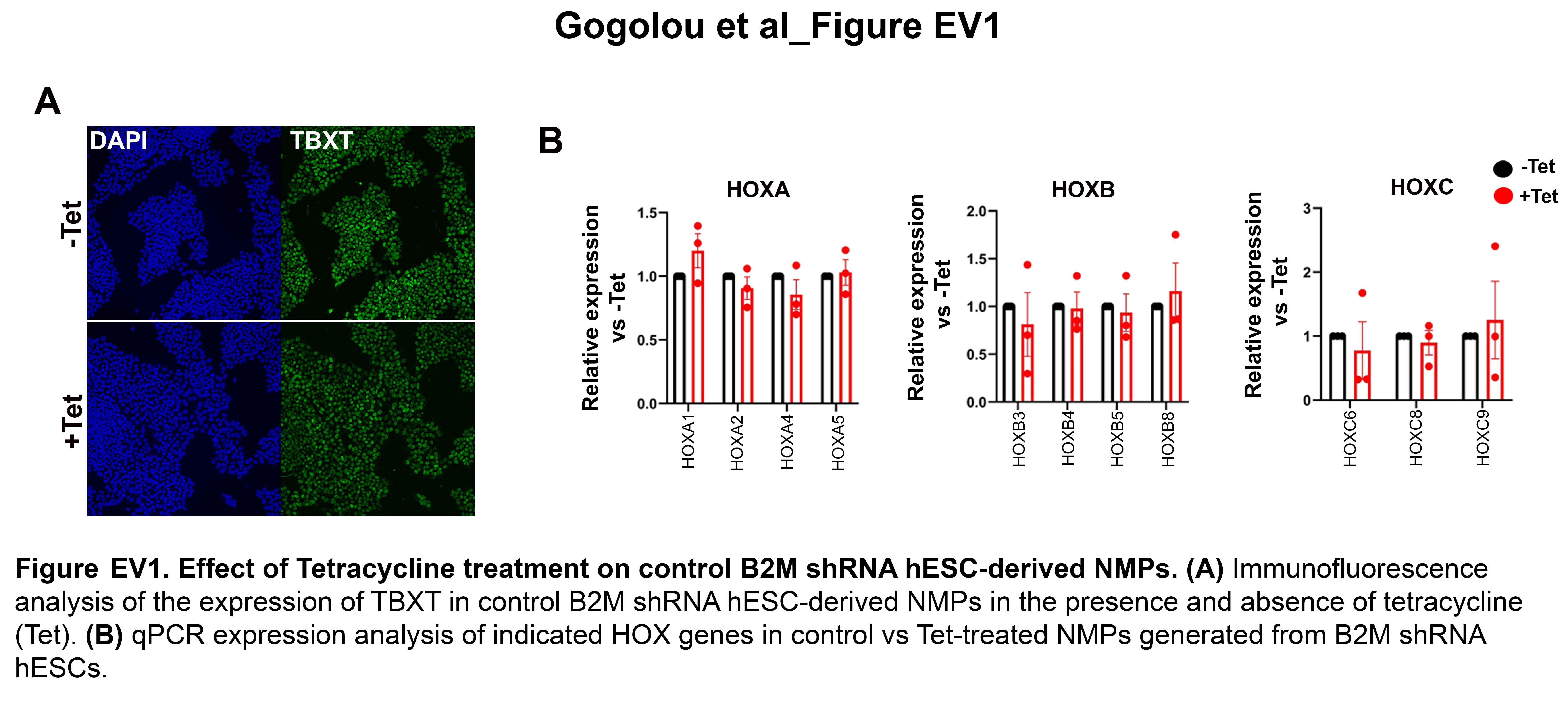

### Fig. EV2

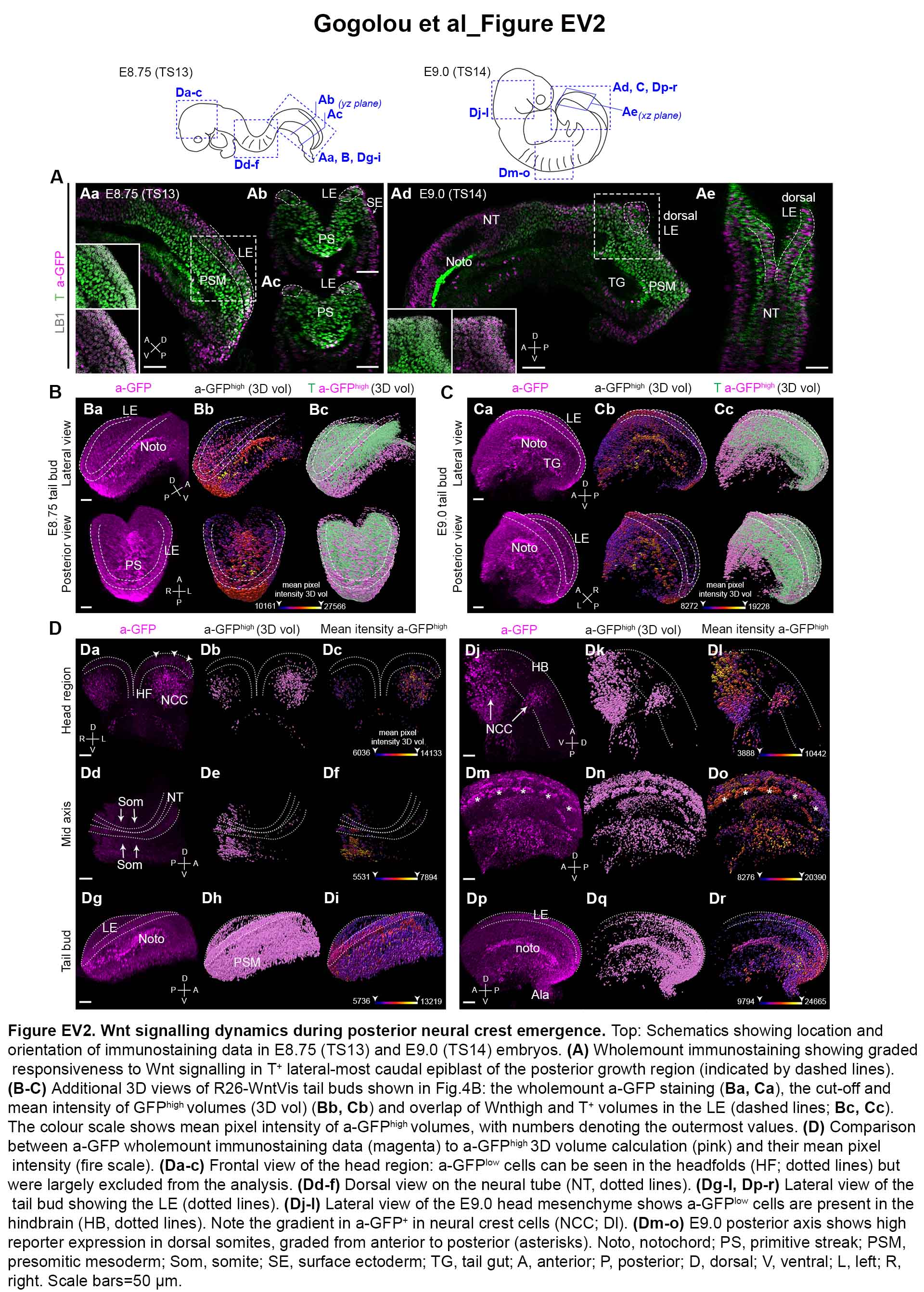

### Fig. EV3

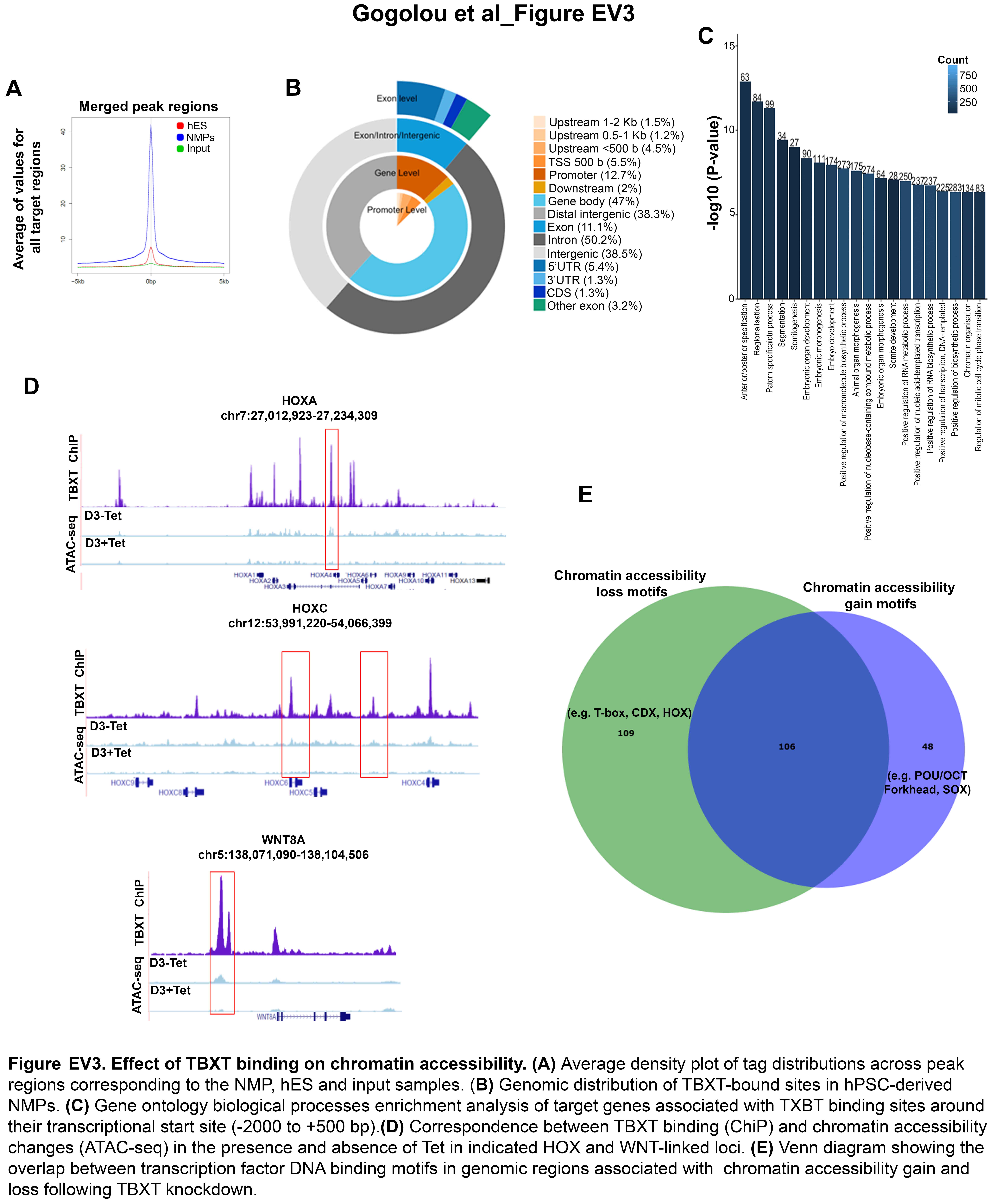
